# Tumour-stroma signaling drives a reversible program with neural-like features in cancer-associated fibroblasts

**DOI:** 10.64898/2026.10.07.757165

**Authors:** Myriam Iliana Ibanez-Rios, Maëlle Batardière, Elham Dianati, Guillaume Serwe, Sajanth Kanagasingam, Alexandre Archambault-Marsan, Pierre Thibault, Marc Therrien, Vincent Quoc-Huy Trinh, David J.H.F. Knapp

## Abstract

Pancreatic ductal adenocarcinoma (PDAC) is one of the most aggressive cancers, in part due to a dense and highly plastic tumor microenvironment dominated by cancer-associated fibroblasts (CAFs). In previous work, we identified a CAF population expressing a neural-associated NES+SOX4+NOTCH1+ program that increased with disease progression. The signals driving this state and the mechanisms regulating it remained unclear. Here, we investigated how tumor-derived signals activate this program and whether it represents a stable fibroblast identity or a reversible state. Using spatial transcriptomic data spanning normal pancreas to invasive carcinoma, we identified EGF, FGF2 and BDNF as candidate tumor-to-fibroblast signals associated with disease progression. Functional experiments in patient-derived CAFs and human fibroblast cell lines showed that EGF and FGF2 are sufficient to induce the NES+SOX4+NOTCH1+ program. Growth factor withdrawal reversed the NES+SOX4+NOTCH1+ program at both the protein and transcriptional levels, indicating that this state is dynamic and environment dependent. Single-cell transcriptomic analysis further revealed a multi-phase response involving early loss of fibroblast-associated features, transient activation of neural-associated genes, and a later stress-associated state. Regulatory analyses identified ETS-family transcription factors as key regulators of the program, while phosphoproteomic profiling and inhibitor studies showed that MAPK/ERK signaling plays a central role in its transcriptional activation. Finally, pan-cancer single-cell analysis showed that this state is enriched in CAFs and increases from normal to tumor tissue across multiple cancer types. Together, these findings define a reversible, growth factor-driven CAF program and identify signaling mechanisms that may provide opportunities to selectively modulate tumor-supportive fibroblast states.

## INTRODUCTION

Pancreatic ductal adenocarcinoma (PDAC) remains one of the most lethal solid tumors, with a five-year survival rate below 12% despite decades of therapeutic advances (1,2). A central contributor to this resistance is the tumor microenvironment (TME), which actively contributes to therapeutic outcome (3,4) . This is exemplified by the dense desmoplastic stroma, composed mainly of cancer associated fibroblasts (CAFs), which physically restricts drug delivery and immune infiltration while actively signaling to support tumor growth (5,6). In healthy tissue, fibroblasts are the main architects of the extracellular matrix (ECM), maintaining tissue structure and homeostasis under normal conditions (7). Upon injury or other stress signals fibroblasts become activated, undergoing transcriptional and epigenetic reprogramming that shifts them toward a proliferative, matrix remodeling phenotype capable of driving wound repair. This activation program is normally transient, resolving once tissue integrity is restored (8).In cancer, however, this same wound-healing response is co-opted and sustained, giving rise to CAFs. CAFs have classically been divided into three subtypes: myofibroblastic CAFs (myCAFs) which express α-SMA (alpha smooth muscle actin) and drive matrix deposition and contractility, inflammatory CAFs (iCAFs) which secrete cytokines and chemokines, and antigen-presenting CAFs (apCAFs) which can modulate adaptative immune responses (9,10). Together, these subtypes exert opposing, context-dependent effects on tumor progression, ranging from tumor-restraining to tumor-promoting. Recent single-cell studies, however, have revealed that this three-way classification captures only part of a much broader spectrum of CAF state (11) . This expanding complexity has made CAFs a difficult therapeutic target, as strategies aimed at broadly depleting or inhibiting CAFs have often produced contradictory outcomes, in some cases accelerating rather than restraining tumor progression (12,13).

Critically, CAF subtype identities are reversible states shaped by the local signaling environment (14–18). The CAF phenotype can therefore be better understood as the output fibroblast-environment interactions than as a marker of committed lineage (19). This plasticity extends even to phenotypes long treated as stable. For example, the canonical myofibroblastic identity, once considered a fixed contractile state, is actively maintained by specific transcriptional switches rather than locked in by terminal differentiation (14). This same plasticity likely underlies the opposing effects described above, as cells shift along a continuum of states rather than occupying fixed, functionally committed identities. Together, these findings reframe the central question in CAF biology away from subtype definition and towards understanding the environmental cues controlling their active program.

CAF plasticity is driven by a dynamic interplay between upstream microenvironmental cues and core intracellular signaling pathways. In the classical model, TGF-β signaling promotes fibroblasts toward a myofibroblastic (myCAF) state, whereas IL-1/STAT2 signaling induces iCAFs. However, central signaling hubs such as PI3K/AKT/mTOR and MAPK/ERK can be activated by distinct upstream cues and integrate these signals to regulate different CAF states and functional programs (20,21). Furthermore, EGFR/ERBB2 activation by the MAPK pathway can drive myCAFs toward a more invasive state (22), demonstrating that the same core signaling pathway can give rise to distinct functional phenotypes even within a defined CAF subtype. This suggests that fibroblast can activate distinct, signal-specific gene programs, each with its own functional consequences, that cut across the classical subtype framework. For example, TGF-β signaling drives a subset of myCAFs to express LRRC15, a program associated with resistance to immune checkpoint blockade across multiple cancer subtypes (23,24). In contrast, blocking TGF-β uncovers a distinct, interferon-licensed fibroblast state with its own specialized immunomodulatory functions (25), highlighting the profound plasticity of these populations.

In our related work on intraductal papillary mucinous neoplasms (IPMN), we identified a subepithelial fibroblast population mainly composed of myCAFs that co-express *NES*, *SOX4* and *NOTCH1*, genes typically associated with neural cell biology (26). Interestingly, the proportion of this CAF population increased with tumor progression (26). Furthermore, we demonstrated that disrupting NOTCH signaling in co-cultures of tumor cell lines and-patient-derived CAFs impairs tumor organoid growth and expansion, underscoring the functional significance of this crosstalk (26) . However, the upstream signals that drive fibroblast into this state, as well as the precise molecular mechanism regulating it, remain unknow.

Here, we set out to identify the tumor-derived signals responsible for inducing the NES+SOX4+NOTCH1+ program and to define the molecular architecture underlying its activation. We hypothesized that specific tumor-secreted growth factors are sufficient to drive fibroblast through this program, and that it represents a reversible, environment-driven program rather than a committed lineage fate. By integrating spatial transcriptomics from the tumor-fibroblast interface from our related work with functional validation, we identified EGF and FGF2 as key tumor-derived drivers and showed that these two growth factors alone are sufficient to activate the program in both patient-derived CAFs and human fibroblast cell lines. We then identified a multi-phase program activation which occurs during phenotype activation using single-cell transcriptomics, and phosphoproteomic profiling and inhibitor cultures to identify necessary upstream signaling drivers. Importantly, we show that this program is fully reversible upon growth factor withdrawal and broadly enriched in CAFs across multiple human cancer types. Altogether, these findings uncover the signaling architecture of NES+SOX4+NOTCH1+ CAFs while establishing that their state is largely environment-dependent, pointing toward alternative strategies for modulating these cells in the TME.

## RESULTS

### Tumor-derived growth factors induce a NOTCH-dependent NES+SOX4+NOTCH1+ phenotype in myCAFs

To characterize how epithelial cells communicate with fibroblasts across IPMN progression, we performed ligand-receptor interaction analysis on spatial transcriptomic data (Xenium *in Situ*) from 18 patients (32 samples covering normal, acinar, low-grade, high-grade and invasive stages). This analysis was restricted to a recurrent grade-associated tissue niche enriched for myCAFs and tumor cells (topic 6) (26). Within this niche, tumor-to-fibroblast signaling at the invasive stage was extensive and involved a broad repertoire of significant ligands spanning growth factors, chemokines, immune-modulatory molecules, and matrix regulators (**Fig. 1A, Supplementary Fig. 1A)**. Notably, interaction strengths for the neural-associated ligands epidermal growth factor (EGF) and brain-derived neurotrophic factor (BDNF) increased progressively from low-grade through invasive carcinoma, whereas fibroblast growth factor 2 (FGF2) was strongly enriched from the low-grade stage onward (**Fig. 1B**). No significant tumor-to-fibroblast interactions were detected in normal tissue (n=1 topic 6 sample, precluding meta-analytic testing) or in acinar tissue (n=6, adequately sampled but no significant hits for any ligand in this panel). Together, these results position tumor-derived EGF, FGF2 and BDNF as the strongest candidates for driving our observed NES+SOX4+NOTCH1+ phenotype (26) in tumour-adjacent fibroblast, with signal strength increasing specifically as disease progresses toward invasive carcinoma.

**Figure 1.**
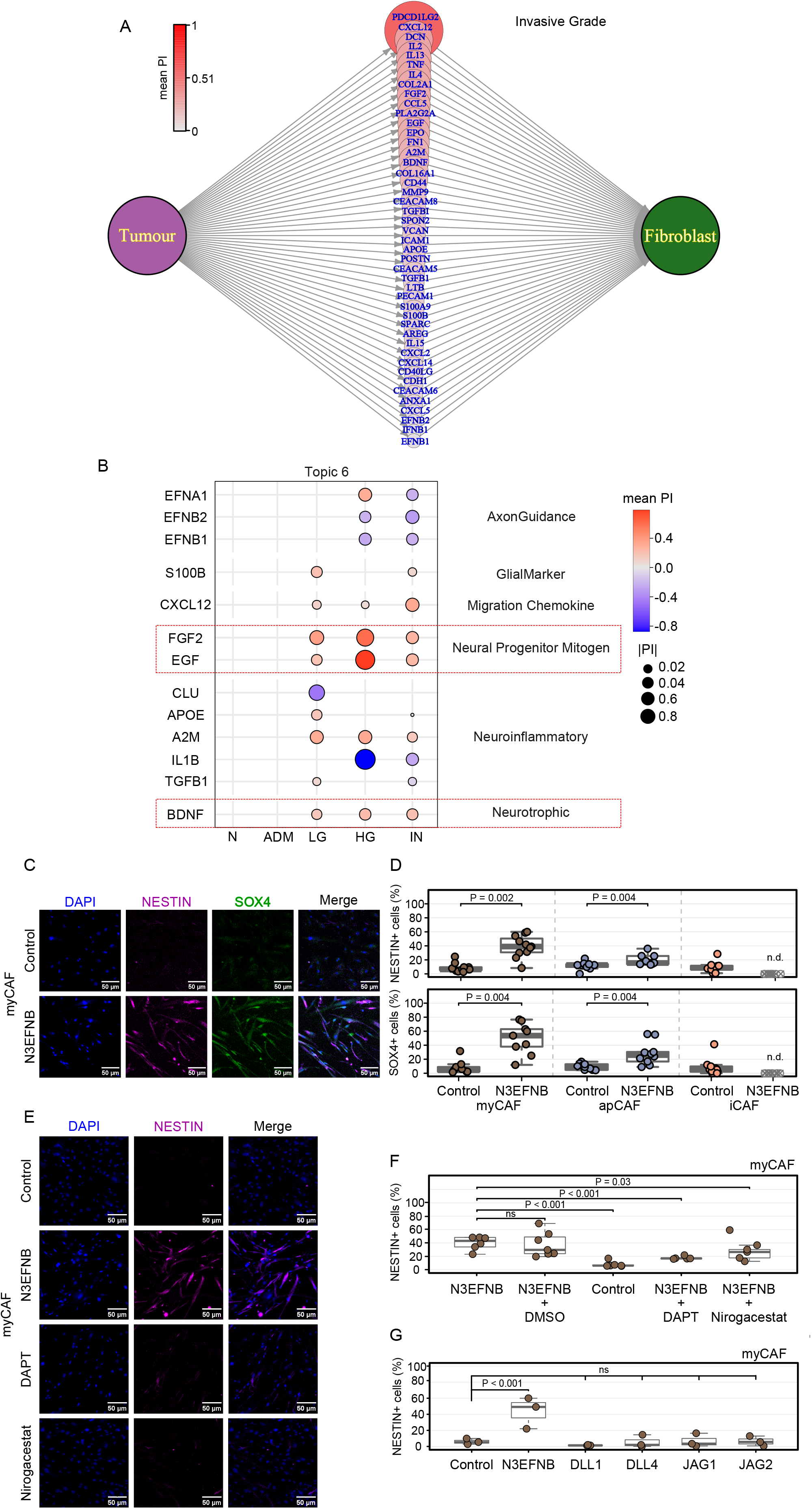
Tumor-derived growth factors activate NES+SOX4+NOTCH1+ phenotype in myCAFs. **A)** Ligands with significant predicted tumor-to-fibroblast interactions at the invasive stage (FDR<0.05, negative interactions are not shown) within the grade-associated topic 6 identified by Xenium spatial transcriptomics (26). **B)** Neural-associated ligands with their fibroblast receptors across histological grades. Only interactions reaching FDR<0.05 are displayed. A blank cell indicates either a non-significant or an untested interaction. Grade-level meta-analysis required at least three contributing samples, so grades below this threshold (n=1) could not be tested. **C)** Representative immunofluorescence images of patient-derived myCAFs after 5 days in control or N3EFNB medium (FGF2, EGF, BDNF and NT3), stained for NESTIN (magenta), SOX4 (green) and DAPI (blue). Scale bar, 50 µm. **D)** Percentage of NESTIN+ and SOX4+ myCAFs and apCAFs after 5 days in control medium or N3EFNB from (**C** and **Supplementary** Fig.1B). Marker positivity was defined relative to within-plate negative controls (see Methods). Each dot represents an experimental replicate performed on a different day (n=9-11 replicates per CAF cell line). iCAFs did not survive N3EFNB treatment and were not quantified (n.d.). Each CAF cell line was analysed separately. Two-sided paired Wilcoxon signed-rank test per CAF cell line, with Benjamini-Hochberg correction across lines. **E)** Representative immunofluorescence images of patient-derived myCAFs after 5 days in control medium, N3EFNB medium with vehicle (0.1% DMSO), DAPT (20 µM) or nirogacestat (10 µM), stained for NESTIN (magenta) and DAPI (blue). Scale bar, 50 µm. **F)** Percentage of NESTIN+ myCAFs from (**E**). Control medium and N3EFNB served as negative and positive controls (n=5-7 experimental replicates). Linear mixed-effects model with biological replicate as a random intercept, followed by Dunnett’s test versus N3EFNB + vehicle. **G)** Percentage of NESTIN+ myCAFs cultured for 5 days in N3 medium (Neurobasal medium with N-2 and B-27 supplements) on plate-bound recombinant DLL1(15 µg/mL), DLL4 (15 µg/mL), JAG1 (15 µg/mL) or JAG2 (5 µg/mL), or uncoated wells in N3 (negative control) or N3EFNB (positive control). n=3 biological replicates. Linear mixed-effects model followed by Dunnett’s test versus N3 control.

Given their significant predicted tumor-to-fibroblast signaling across multiple grades, we selected EGF, FGF2 and BDNF for functional validation. We also included neurotrophin-3 (NT3), which was not present in the Xenium gene panel, but which has an established role in both neural progenitor differentiation and pancreatic cancer biology (27). To test whether tumor-derived growth factors are sufficient to activate the NES+SOX4+NOTCH1*+* phenotype in CAFs, we cultured pancreatic-tumour-derived CAFs of different subtypes (myCAFs, iCAFs and apCAFs) in control or neurobasal medium containing EGF, FGF2, BDNF and NT3 (N3EFNB medium) for five days. Following treatment, myCAFs robustly upregulated both NESTIN and SOX4, compared to near baseline labels in uninduced controls (**Fig. 1C-D**). By contrast, apCAFs exhibited only modest activation, while iCAFs failed to survive the treatment conditions, precluding assessment on their responsiveness (**Fig. 1D and Supplementary Fig. 1B**). This myCAF-associated effect is consistent with the spatial enrichment of *NES* and *SOX4* transcripts in myofibroblastic stroma documented we previously documented (26).

We next asked which downstream pathway mediates this induction. NOTCH1 is one of the defining markers of the NES+SOX4+NOTCH1+ phenotype identified in our related study, where NOTCH inhibition reduced tumor growth in tumor cells-myCAF cells co-culture (26). We therefore tested whether NOTCH signaling is required for the induction of the phenotype itself. myCAFs were cultured in N3EFNB medium for 5 days in the presence or absence of gamma-secretase inhibitors DAPT or nirogacestat, which inhibit the cleavage of NOTCH receptors inside the cell membrane. Interestingly, both inhibitors significantly reduced the percentage of NESTIN+ cells relative to the N3EFNB condition alone (**Fig. 1E-F**). To determine whether direct NOTCH ligand engagement is sufficient to induce the NESTIN+ phenotype independent of growth factor signaling, myCAFs were plated onto wells coated with recombinant DLL1, DLL4, JAG1 or JAG2 (28) , alongside control untreated and N3EFNB treated cells as negative and positive controls respectively. As before, N3EFNB robustly induced NESTIN relative to control, whereas none of the four NOTCH ligands increased the percentage of NESTIN+ above control levels (**Fig.1G**). These results indicate that NOTCH signaling is necessary but not sufficient to drive the NESTIN phenotype in myCAFs.

### EGF and FGF2 are partially redundant inducers of a reversible NES+SOX4+NOTCH1+ fibroblast phenotype

While primary patient-derived CAF lines confirmed that tumor-derived growth factors are sufficient to activate the NES+SOX4+NOTCH1+ phenotype in myCAFs, their rapid entry into replicative senescence and subsequent phenotypic drift *in vitro* limit their use for detailed mechanistic studies(10,29,30). To circumvent this restriction and establish and scalable model, we used standard human fibroblast cell lines (BJ, MRC5 and IMR90). Following N3EFNB treatment these induced strong activation of NESTIN, SOX4, and NOTCH, similar to the primary myCAFs (**Fig. 2A-B, Supplementary Fig. 2A**). Additionally, we assessed TUJ1 (βIII-tubulin), a neuronal marker associated with neural differentiation which was also induced under N3EFNB stimulation (**Supplementary Fig.2A-C**).

**Figure 2.**
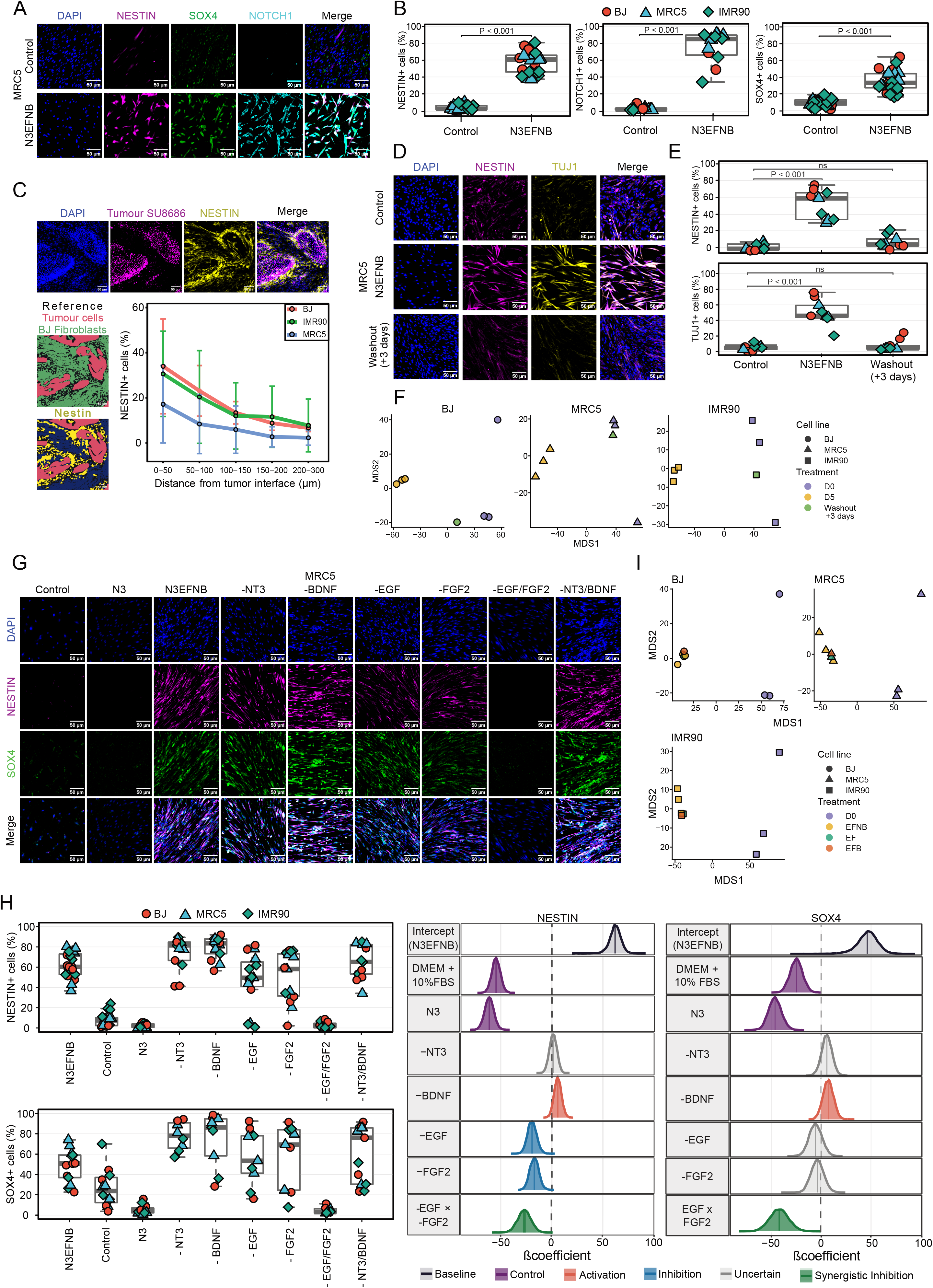
EGF and FGF2 are partially redundant inducers of a reversible NES+SOX4+NOTCH1+ fibroblast phenotype. **A)** Representative immunofluorescence images of MRC5 fibroblasts after 5 days in control or N3EFNB medium, stained for NESTIN (magenta), SOX4 (green), NOTCH1 (cyan) and DAPI (blue). Scale bar, 50 µm. **B)** Percentage of NESTIN+, SOX4+ or NOTCH1+ fibroblasts from (**A** and **Supplementary** Fig. 2A). Marker positivity was defined relative to within-plate negative controls (see Methods). Each dot represents one experimental replicate. Linear mixed-effects model with biological replicate and cell line as random intercepts, followed by Dunnetts’s test versus control. **C)** Representative immunofluorescence images of BJ in co-culture with SU8686 (magenta), stained for NESTIN (yellow) and DAPI (blue), and the corresponding segmentation map used for quantification, showing NESTIN+ (yellow) and NESTIN-(blue) BJ fibroblasts and SU8686 cells (pink). Scale bar, 50 µm. Percentage of NESTIN+ fibroblasts according to distance from the nearest tumor cell in 5-day co-cultures of fibroblasts from (**C** and Supplementary Fig. 2D) and SU8686 (5:1 ratio) in growth factor-free medium. Points show the mean and error bars show the SD per cell line (BJ and MRC5, n=4; IMR90, n=3 experimental replicates). Linear regression of NESTIN+ percentage against distance in each well, with slopes analysed by a linear mixed-effects model with experiment as a random intercept. **D)** Representative immunofluorescence images of MRC5 fibroblasts in control medium, after 5 days of N3EFNB treatment, and after 5 days in N3EFNB medium followed by 3 days in control medium (DMEM+10%FBS; washout). Stained for NESTIN (magenta), TUJ1 (yellow) and DAPI (blue). Scale bar, 50 µm. **E)** Percentage of NESTIN+ and TUJ1+ fibroblasts from (**D** and **Supplementary** Fig.2E) (BJ, MRC5 and IMR90; n=3 experimental replicates per cell line). Linear mixed-effects model with replicate and cell line as random intercepts, followed by Dunnett’s test versus control. **F)** Multi-dimensional scaling (MDS) of bulk RNA-sequencing profiles from control (D0; n=3 per cell line), N3EFNB-treated (D5; n=3 per cell line), and washout (n=1 per cell line) samples, based on 2,297 genes differentially expressed between D5 and D0 (|log2FC| > 1, FDR < 0.01; voom-normalized expression, leading log2FC method). **G)** Representative immunofluorescence images of MRC5 fibroblasts in control medium, after 5 days of N3EFNB lacking one factor (−EGF, −FGF2, −BDNF, −NT3) or a pair of factors (−EGF/−FGF2 or −BDNF/−NT3) culture with complete N3EFNB-treatment as positive control and N3 medium as negative control. Stained for NESTIN (magenta), SOX4 (green), and DAPI (blue). Scale bar, 50 µm. **H)** Percentage of NESTIN+ and SOX4+ fibroblasts from (**G** and **Supplementary** Fig. 2F). Forest plots showing the posterior distribution of each effect (difference in the percentage of positive cells relative to N3EFNB) from Bayesian mixed-effects models with cell line and biological replicate as random intercepts. Dark shading indicates the 95% credible interval, and vertical lines indicate posterior medians (n=4-7 NESTIN and n=3-5 SOX4 experimental replicates per cell line per condition). **I)** MDS of bulk RNA-sequencing profiles from control (n=3 per cell line), N3EFNB-treated (n=3 per cell line), EGF/FGF2 (EF; n=1 per cell line), and EGF/FGF2/BDNF (EFB; n=1 per cell line) samples, based on the same DEG set defined in (**F**).

Considering that *NES* is upregulated in myCAFs near the tumor edge (26), we asked whether PDAC tumor cells (SU8686) could induce NESTIN expression in human fibroblast (BJ, MRC5 and IMR90) when co-cultured. In all three fibroblast cell lines, the proportion NESTIN+ fibroblasts was higher immediately adjacent to the tumor interface and progressively declined with increasing distance from the tumor cells (P = 0.037; **Fig. 2C, Supplementary Fig. 2D**). Together, these data suggest that tumor cells induce a spatially restricted NESTIN response in fibroblast, supporting a model of distance-dependent tumor-stromal crosstalk. Given that this phenotype could be induced by both N3EFNB treatment and tumor cell proximity, we next asked whether it represents a stable lineage commitment or a reversible phenotypic state.

To address this, we performed a washout experiment, replacing the N3EFNB treatment after 5 days with standard culture medium for 3 days (DMEM+10%FBS). Upon growth factor withdrawal, NESTIN and TUJ1 positivity dropped to levels indistinguishable from the baseline conditions (**Fig. 2D-E, Supplementary Fig. 2E**). Furthermore, to determine whether this reversibility extends beyond protein expression to the broader transcriptional program, we performed bulk RNA-sequencing on the three fibroblast cell lines (BJ, MRC5 and IMR90) under control, N3EFNB-treated, and washout conditions. Differentially expressed genes (log2FC>1, FDR<0.01) between treated and control samples (n=2,297 genes; **Supplementary Tables: 4**) were used for multidimensional scaling of voom-normalized expression values. This demonstrated clear separation of N3EFNB-treated samples from control along the first dimension, while the washout sample clustered closely with controls in each cell line (**Fig. 2F**). Together, these data are consistent with the NES+SOX4+NOTCH1+ program representing a dynamic, growth factor-dependent activation state rather than a permanent lineage fate.

Having established that the NES+SOX4+NOTCH1+ program is reversible, we next aimed to determine which components of the N3EFNB cocktail were required for this activation. We performed a factorial omission experiment in BJ, MRC5 and IMR90 fibroblast, withdrawing FGF2, EGF, BDNF or NT3 from N3EFNB medium, either individually or in pairs (-EGF/-FGF2, - BDNF/-NT3). Combined withdrawal of EGF and FGF2 abolished the program, reducing NESTIN, SOX4 and TUJ1 to levels comparable to growth factor free medium (**Fig. 2G-H, Supplementary Fig. 2F-H**). However, the three markers differed in how they depend in each factor. NESTIN was reduced by the withdrawal of EGF and FGF2 alone (P(effect) > 0.999), though each single omission accounted for only about one third of the full response. A strong negative EGF × FGF2 interaction (P(effect) > 0.999) indicated that either factor sustained most of NESTIN induction. SOX4 showed redundancy as withdrawal of EGF or FGF2 alone had a much weaker effect (P(effect) = 0.87 and 0.82 respectively), whereas their combined removal abolished SOX4 expression through a strong negative interaction (P(effect) > 0.999). In contrast, TUJ1 depended additively on both factors, with comparable reductions upon removal of EGF or FGF2 (P(effect) > 0.999) and no support for an interaction. In contrast, the withdrawal of neurotrophic factors BDNF and NT3 had little effect on the phenotype while EGF and FGF2 in combination (ie. -BDNF/-NT3) were sufficient to induce comparable levels of NESTIN+, SOX4+, and TUJ1+ cells relative to the complete cocktail (**Fig. 2G-H, Supplementary Fig. 2F-H**). We further assessed these conditions by bulk RNA-sequencing, comparing N3EFNB, EGF/FGF2, and EGF/FGF2/BDNF across the three fibroblast cell lines (BJ, MRC5 and IMR90). MDS analysis using the same DEG set identified in the reversibility experiment showed that both EF and EFB remained close to the complete N3EFNB condition and separated from untreated controls (**Fig. 2I, Supplementary Tables: 4)**. Together, these results indicate that EGF and FGF2 are sufficient to induce the NES+SOX4+NOTCH1+ program at both the protein and transcriptional levels.

We next performed dose-response experiments to determine the optimal concentration of EGF and FGF2 for program induction. NESTIN positivity reached a plateau from 5 ng/mL onward for both factors, individually and in combination (P < 0.001 versus control for all conditions ≥ 5 ng/mL; **Supplementary Fig. 2I**). We selected 20 ng/mL for both EGF and FGF2 as a robust concentration within this plateau. Overall, these findings establish the NES+SOX4+NOTCH1+ program as a reversible fibroblast state driven primarily by EGF and FGF2.

### Dynamic transcriptional changes during N3EFNB treatment

To define how the NES+SOX4+NOTCH1+ program develops over time, we performed scRNA-sequencing on BJ, MRC5 and IMR90 fibroblasts at D0, D2 and D5 of N3EFNB treatment. AML5 cells and iPSC-derived neural progenitor cells (NPCs) were included as reference populations and to control for time-associated batch effects (**Fig. 3A**). Cell identities were resolved by genotype deconvolution and confirmed using population-specific markers and targeted genotyping (**Supplementary Fig. 3A-E**). Pseudotime analysis independently recapitulated experimental time in each fibroblast line, with cells progressing continuously from D0 toward D5 rather than separating into discrete transcriptional states (**Fig. 3B, Supplementary Fig. 3F**). Consistent with this trajectory, consensus integrative NMF (ciNMF; k=13) identified three major time-associated programs shared across the fibroblast lines (**Fig. 3C, Supplementary Tables: 5**). A fibroblast/ECM-associated program (Factor 1; *COL1A2, DCN, ZEB2*) was highest at D0 and declined with induction, while a program associated with neuronal cytoskeletal and lipid-biosynthesis program (Factor 4; *TUBB3* (*TUJ1), MSMO1, FDFT1, FDPS*) peaked transiently at D2. At D5, the strongest associated program was enriched for stress and growth-arrest genes (Factor 2; *CDKN1A, GDF15, TIMP1, SERPINE2*). Importantly, the neural progenitor program present in NPCs (Factor 8; *PAXC, FOXG1, SIX3*) was not appreciably induced in fibroblasts, indicating that N3EFNB activates selected neural-associated features without driving cells toward a bona fide neural progenitor identity, consistent with their failure to repopulate a cerebral organoid following transplantation (31) **(Supplementary Figure 4A-B**). Together these results confirm that while the fibroblasts gain a phenotype with many properties of neural progenitors, they do not undergo true lineage conversion. We validated these temporal changes by comparing gene expression over time of *ACTA2*, *NES*, *TUBB3* (*TUJ1)*, *SOX4*, *SOXC, SOXS* with RT-qPCR in MRC5 fibroblasts at D0, D2 and D5 (**Fig. 3D**). *NES*, *TUBB3* (*TUJ1)* and *SOX9* were significantly induced by D2 and remained elevated at D5, whereas *SOX6* and *SOX4* showed delayed induction and were significantly increased only at D5 (**Fig. 3D**). Conversely, the fibroblast marker *ACTA2* was transiently reduced at D2.

**Figure 3.**
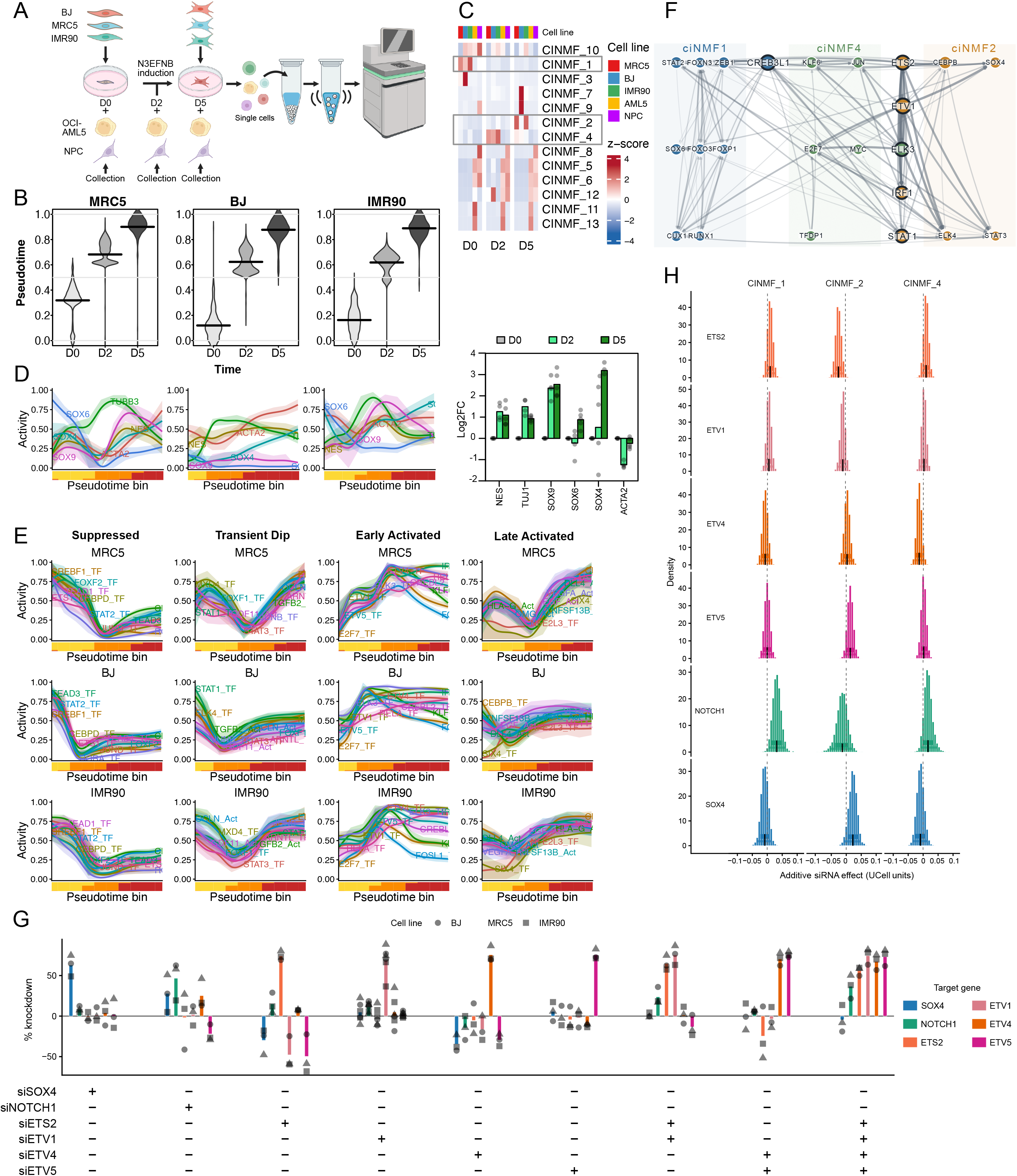
Dynamic transcriptional changes during N3EFNB treatment reveal ETS-family control of the NES+SOX4+NOTCH1+ program in fibroblasts. A) Schematic of the scRNA-sequencing experiment. BJ, MRC5 and IMR90 fibroblasts were collected at D0, D2 and D5 of N3EFNB treatment; OCI-AML5 cells and iPSC-derived NPCs were included as reference populations. B) Pseudotime distributions of MRC5, BJ and IMR90 fibroblasts at D0, D2 and D5, showing continuous progression from D0 to D5 during N3EFNB treatment. C) Heatmap of ciNMF program activity across D0, D2 and D5 fibroblasts and the OCI-AML5 and NPC reference populations. Values represent row-scaled z-scores. The ciNMF programs associated with the D0, D2 and D5 fibroblast states are highlighted. D) Pseudotime-resolved marker gene expression in MRC5, BJ and IMR90 fibroblasts (left) and RT-qPCR expression of the same genes in MRC5 fibroblasts at D0, D2 and D5, shown as log2 fold-change relative to D0 (right). For the pseudotime resolved markers both here and in (E), the median and interquartile range (25th–75th percentile) were calculated within each bin and smoothing spline fit to the bin means to generate a continuous trajectory, with the interquartile range displayed as a shaded ribbon. A stacked bar beneath each trajectory shows the proportion of cells from each experimental timepoint (D0, D2, and D5) represented within each bin (D0=yellow, D2=orange, D5=red). E) Pseudotime-resolved transcription factor regulon and inferred ligand activities in MRC5, BJ and IMR90 fibroblasts, showing distinct temporal regulatory patterns during N3EFNB treatment. F) Regulatory network linking the top SCENIC-inferred transcription factor regulons to genes associated with the D0-, D2- and D5-associated ciNMF programs (ciNMF-1, ciNMF-4 and ciNMF-2, respectively). G) Percentage knockdown of *SOX4, NOTCH1, ETS2, ETV1, ETV4* and *ETV5* measured by bulk RNA-sequencing following individual or combinatorial siRNA treatment in BJ, MRC5 and IMR90 fibroblasts (n = 1 per condition per cell line). H) Additive effects of individual siRNA knockdowns on D0-, D5- and D2-associated ciNMF program activity (ciNMF-1, ciNMF-2 and ciNMF-4, respectively), quantified from bulk RNA-sequencing and expressed in UCell units. Posterior distributions were estimated using Bayesian models.

To better understand the control of these cell programs, we next examined the activity of transcription factor regulons and inferred ligand activities along the temporal trajectory. Clustering based on pseudotime-resolved activity identified reproducible temporal patterns across the fibroblast lines (**Fig. 3E, Supplementary Figure 4C-D).** Fibroblast-associated regulators, including CREB3L1, JUND, SREBF1, CEBPD, TEAD3 and FOXF2, were rapidly suppressed. This was followed by early activation of regulators including ETV4, ETV1, KLF6, ELK3, IRF1, ETV5 and FOSL1, together with a transient cluster containing STAT3, ELK4, STAT1, JUNB and MXD4. Interestingly, the late-activated cluster was dominated by inferred ligand activities, including DLL4, HLA-G, TNFSF13B, OMG, VEGFA and EGF. Together, these results support a continuous, multi-phase transcriptional response to N3EFNB, characterized by early loss of fibroblast identity, transient activation of MAPK-associated TFs and neural-associated genes, and a later shift toward a growth arrested secretory associated state.

### Regulatory control of NES+SOX4+NOTCH1+ program in fibroblasts is driven by ETS-family transcription factors

To identify candidate transcriptional regulators underlying the shared D0→D2→D5 trajectory common to all three fibroblast cell lines (Factors 1, 4, and 2 respectively), we intersected each program’s top loading genes with SCENIC-inferred regulons (**Supplementary Tables: 6**). We examined not only which regulators covered each program, but how concentrated or distributed that regulatory control was, using marginal coverage (new genes contributed beyond higher-ranked regulators) and pairwise containment (the fraction of a regulator’s covered genes also covered by another) among the top 10 regulators per program (**Fig. 3F**). Several regulators showed high target coverage of more than one program: CREB3L1 was shared between the D0 (Factor 1) and D2 (Factor 4) programs, while ETS2, ETV1, and ELK3 were shared between the D2 (Factor 4) and D5 (Factor 2) programs, consistent with the transition through the shared trajectory being driven by an overlapping, partially continuous regulatory cascade rather than entirely distinct transcription factor sets at each stage. Within the D5 program specifically, ETS2 and ETV1 showed near-complete target overlap, indicating redundant rather than independent regulatory contributions at this stage.

In order to determine the functional implications of some of the core factors suggested by the adjacency matrix to control the observed time-dependent transcriptional programs, we performed siRNA knockdown in BJ, MRC5, and IMR90 fibroblasts and measured the impacts on both targeted genes as well as the time-dependent programs. Single knockdowns achieved robust reduction of their intended target, and combination knockdowns achieved comparable efficiency across all targeted genes simultaneously, including the triple-plus-*ETS2* combination (**Fig. 3G**). Beyond the intended targets, several knockdowns were accompanied by compensatory changes in other pathway members’ transcript levels. Notably, knockdown of *ETS2, ETV1*, or *ETV4* individually was consistently associated with increased expression of the other *ETS*-family/AP-1-adjacent factors (e.g., siETS2 increased *ETV1* and *ETV5* transcript levels, siETV4 increased *SOX4*, *ETV1*, and *ETV5*), while siNOTCH1 was associated with reduced *SOX4* and *ETV4* transcript levels. These secondary changes are consistent with cross-regulatory relationships among the candidate regulators identified above.

To test whether the candidate regulators identified above are functionally required for ciNMF program activity, rather than merely correlated with it, we used the additive Bayesian models to estimate each siRNA’s independent effect on each of the time-associated gene programs (ciNMF_1, ciNMF_4, ciNMF_2). Knockdown of *NOTCH1* significantly increased the initial fibroblast state-associated ciNMF_1 program score along with the early activated program (P(effect) = 0.991 & 0.955 respectively; **Fig. 3H, Supplementary Tables: 7**). This pattern suggests that *NOTCH1* is likely involved in suppression of these programs. Knockdown of *ETV4* appeared to have a positive regulatory effect on the early program ciNMF_4 (P(effect) = 0.947) based on the negative effect of knocking it down, consistent with its membership in the Early Activated cluster identified above. Knockdown of *ETS2* and to a lesser extent *ETV1* were associated with a decrease in the late program, ciNMF_2 (P(effect) = 0.991 & 0.879 respectively, **Fig. 3H, Supplementary Tables: 7**) suggesting they’re involved in promoting it. This is consistent with their regulatory links to ciNMF_2 associated genes from the SCENIC adjacency matrix. Interestingly, both *ETV5* and *SOX4* knockdown were associated with a weak *increase*, in ciNMF_2 (P(effect) = 0.828 & 0.945 respectively), suggesting these genes may actually restrain the late activation program.

### MAPK/ERK and PI3K/AKT signaling differentially regulate the NES+SOX4+NOTCH1+ program

We next aimed to identify the intracellular signaling pathways involved in the NES+SOX4+NOTCH1+ program activation. We performed time-course phosphoproteomic profiling on MRC5 fibroblasts following combined EGF and FGF2 stimulation (0, 10, 30 and 120 mins). KEGG pathway-enrichment analysis identified several pathways associated with growth factor signaling, including ErbB signaling (P = 4.55 × 10⁻⁷) and MAPK signaling (P = 9.73 × 10⁻⁴, **Fig. 4A**). Notably, we detected phosphorylation changes in several components of MAPK/ERK cascade, including EGFR, RAF1, MAP2K1 (MEK1) and MAPK1 (ERK2), as well as the downstream effectors RPS6KA1 and RPS6KA3. Changes were also detected in PI3K-associated proteins, including PIK3R2 and MTOR. Based on these findings, we next investigated the contribution of MAPK/ERK and PI3K/AKT signaling to the *NES+SOX4+NOTCH1+* program.

**Figure 4.**
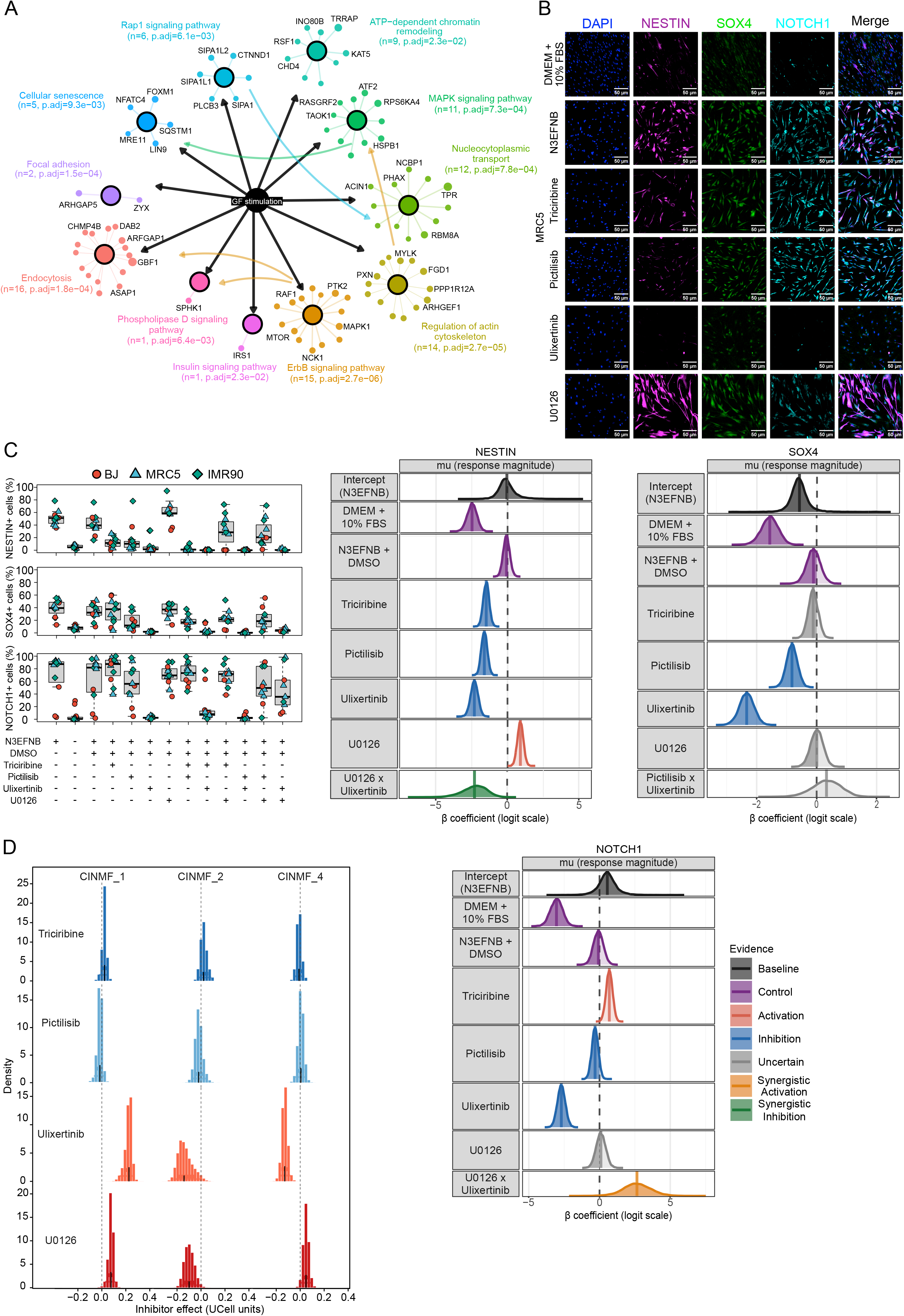
MAPK/ERK and PI3K/AKT signaling differentially regulate the NES+SOX4+NOTCH1+ program. **A)** KEGG pathway-enrichment network of significantly regulated phosphoproteins in MRC5 fibroblasts following combined EGF and FGF2 stimulation. Protein nodes are connected to their associated enriched pathways, with node size representing −log₁₀(P-value) of the most significant phosphosite. Curved arrows indicate schematic relationships between enriched pathways supported by KEGG pathway maps and were not inferred from the phosphoproteomic data, n indicates the number of phosphoproteins displayed for each pathway after assignment to their most significantly enriched pathway (42,43). **B)** Representative immunofluorescence images of MRC5 fibroblasts cultured for 5 days in N3EFNB medium with vehicle or pathway inhibitors, stained for NESTIN (magenta), SOX4 (green), NOTCH1 (cyan) and DAPI (blue). Scale bar, 50 µm. **C)** Percentage of NESTIN+, SOX4+ and NOTCH1+ fibroblasts from (**C** and **Supplementary** Fig. 6A) following treatment with pathway inhibitors, alone or in combination. Box plots show the percentage of positive cells and forest plots show the posterior distribution of each inhibitor effect relative to N3EFNB + vehicle. Marker positivity was defined relative to within-plate negative and positive controls (see Methods). Bayesian mixed-effects models included biological replicate and cell line as random intercepts. Dark shading indicates the 95% credible interval, and vertical lines indicate posterior medians (BJ, n = 3; MRC5 and IMR90, n = 4 experimental replicates). **D)** Activity of the D0-, D2- and D5-associated ciNMF programs (ciNMF-1, ciNMF-4 and ciNMF-2, respectively) in bulk RNA-sequencing samples from BJ, MRC5 and IMR90 fibroblasts cultured for 5 days in N3EFNB medium with AKT (Triciribine), PI3K (Pictilisib), MEK (U0126) or ERK (Ulixertinib) inhibitors. Program activity was quantified by scoring the expression of genes associated with each ciNMF program. Inhibitor effects represent changes relative to matched N3EFNB + DMSO controls. Posterior distributions were estimated using Bayesian mixed-effects models with cell line as a random intercept.

We first asked whether these inhibitors impacted the activation of the NES+SOX4+NOTCH1+ phenotype. As expected, NESTIN, SOX4 and NOTCH1 were each significantly reduced by Ulixertinib (P(effect) > 0.999). Interestingly, U0126 did not reduce SOX4 or NOTCH1 and increased NESTIN (P(effect) > 0.999; **Fig. 4B-C, Supplementary Fig. 6A**) despite both U0126 and Ulixertinib markedly reducing ERK and RSK phosphorylation by western blot (**Supplementary Fig. 6B**). Pictilisib reduced NESTIN and SOX4 (P(effect) > 0.999) but had a weaker effect on NOTCH1 (P(effect) = 0.917) while Triciribine reduced NESTIN (P(effect) > 0.999) with a limited effect on SOX4 (P(effect) = 0.765) and actually increased NOTCH1 (P(effect) = 0.999; **Fig. 4B-C, Supplementary Fig. 6A**). Interestingly, TUJ1 responded differently, where Pictilisib and Triciribine reduced it, while Ulixertinib and U0126 increased it (all P(effect) > 0.999; **Supplementary Fig. 6C-D**). These results highlight that MAPK and AKT signaling play a role in the phenotype activation, however, as we saw in the program membership, these phenotypic markers are associated with different sub-programs and are terminal leaves, thus their regulation is more complex.

Given that the protein-level phenotype reflects the activity of multiple phases of the activation state of the NES+SOX4+NOTCH1+ program, we performed bulk RNA-sequencing to provide a complete view of the effects of inhibiting these pathways on program activity. Normal fibroblasts were exposed to N3EFNB for 5 days in the presence of inhibitors or vehicle controls, then scored for the ciNMF programs identified from the scRNA-sequencing time course (**Fig. 4D**). Direct ERK inhibition with Ulixertinib produced the strongest transcriptional shift, decreasing the D2- and D5-associated programs (ciNMF-4, P(effect) > 0.999; ciNMF-2, P(effect) = 0.969, respectively), while increasing the D0-associated program (ciNMF-1, P(effect) > 0.999). U0126 produced a smaller shift in the same direction for D0 and D5 (ciNMF-1, P(effect) = 0.993; ciNMF-2, P(effect) = 0.987) but increased rather than decreased D2 (ciNMF-4, P(effect) = 0.979). Pictilisib and Triciribine had comparatively limited effects, with no program-level shift exceeding P(effect) = 0.884 in either direction. These results indicate that direct ERK inhibition shifts the transcriptional state away from the NES+SOX4+NOTCH1+ program and toward the baseline-associated state. Despite a clear effect on NESTIN in these lines (**Supplementary Fig. 7A-C**), as we had observed in myCAFs, DAPT and nirogacestat both had relatively limited and relatively inconsistent effects on overall program activity (largest shift in any program: ciNMF-4, nirogacestat, median +0.041 UCell units, P(effect) = 0.968; **Supplementary Fig. 7D)**. Overall, these data confirm that NOTCH1 and PI3K/AKT signaling primarily modulates individual marker protein levels, whereas MAPK/ERK signaling is more closely linked to regulation of the broader NES+SOX4+NOTCH1+ transcriptional program.

### The NES+SOX4+NOTCH1+ program is enriched in cancer-associated fibroblasts across cancer types

Having defined the signaling pathways regulating the NES+SOX4+NOTCH1+ program *in vitro*, we next asked whether this transcriptional state could also be identified across human cancer types. To evaluate the presence of this program, we reanalyzed a published pan-cancer single-cell RNA-seq atlas ((11); GEO: GSE210347), comprising 855,271 cells from 234 samples across 11 tumor types and normal, tumor-adjacent, and tumor tissues (**Fig. 5A**). We first asked whether the early and late transcriptional programs (ciNMF-4 and ciNMF-2, respectively) were restricted to fibroblasts or broadly expressed across the tumor microenvironment. Scoring these programs across all major cell populations showed that D5 program activity was concentrated almost entirely within the fibroblast compartment (**Fig. 5B-C**). In contrast, the D2 program was distributed more broadly across multiple cell types (**Fig. 5B-C**), indicating that the late D5 state is more fibroblast-restricted than the earlier transcriptional response. We next asked whether late program activity distinguished CAFs from normal fibroblasts (nlFBs) within this compartment. After subsetting the fibroblast population only (n=104,804 cells) and re-scoring the D5 program, we found that activity was strongly enriched in CAFs compared with nlFBs (**Fig. 5D**). To further identify cells resembling the NES+SOX4+NOTCH1+ phenotype characterized *in vitro*, we additionally scored a three-gene marker signature comprising *NES, SOX4,* and *NOTCH1*. Late program activity and *NES/SOX4/NOTCH1* positivity co-occurred significantly more often than expected by chance, with D5 program-positive fibroblasts having approximately twice the odds of also being *NES/SOX4/NOTCH1* marker-positive (OR = 2.00, 95% CI 1.85–2.17; mixed-effects logistic regression, P < 0.001). This association was observed across samples and tumor types, with an OR > 1 in 158 of 178 samples (median OR = 2.24) and identified a late program-positive + *NES/SOX4/NOTCH1*-positive population representing 9.2% of all fibroblasts.

**Figure 5.**
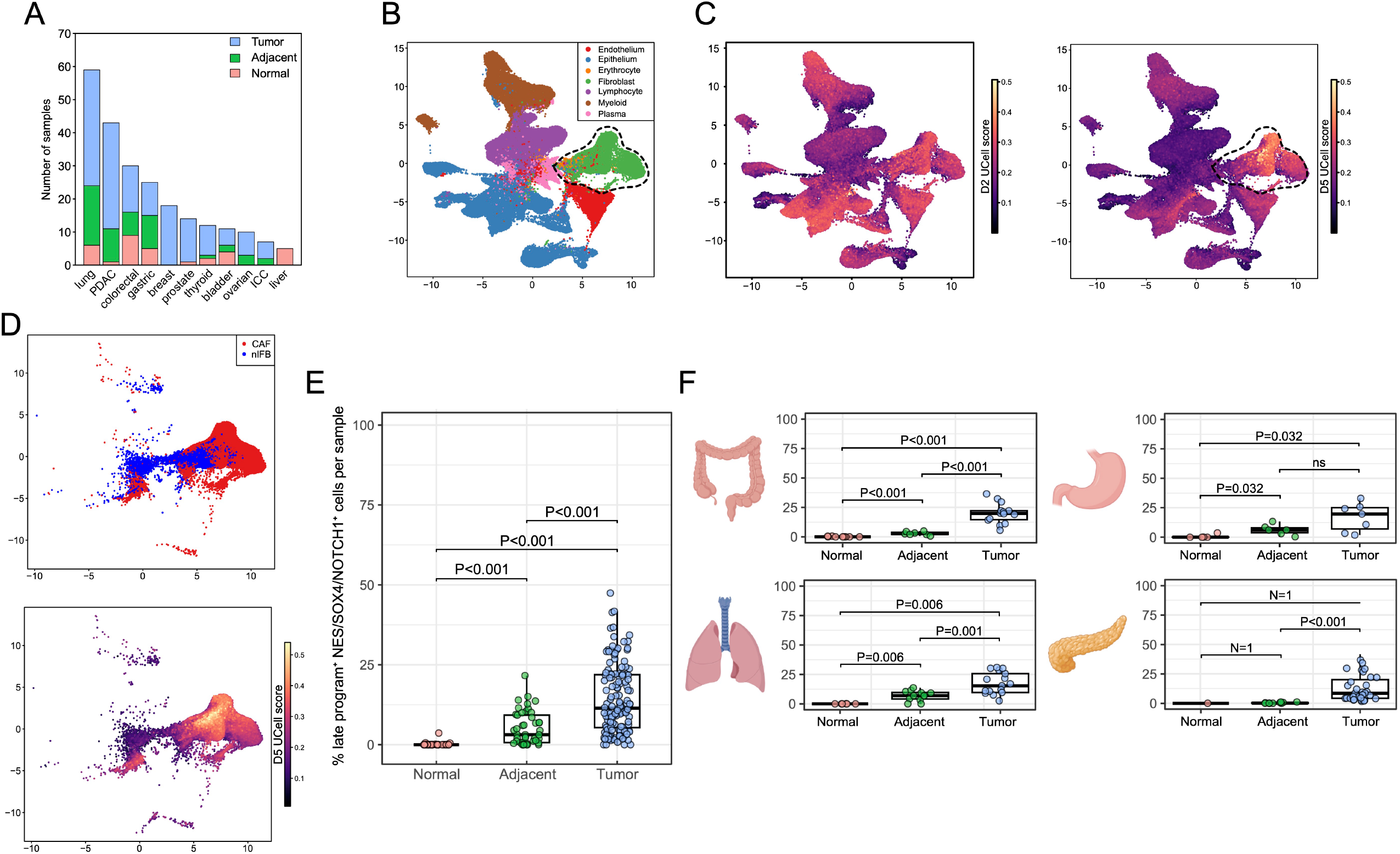
The NES+SOX4+NOTCH1+ program is enriched in cancer-associated fibroblasts across cancer types. **A)** Number of samples from each cancer type stratified by tissue state (normal, tumor-adjacent, and tumor) from pan-cancer single-cell RNA-sequencing atlas re-analysis (11) (GEO: GSE210347). **B)** UMAP of 855,271 cells colored by major cell type (left). The fibroblast compartment is indicated by the dashed outline. **C)** Same UMAP colored by D2- and D5-associated program score (ciNMF-4 and ciNMF-2, respectively). **D)** UMAP of the fibroblast compartment (n = 104,804 cells) colored by cancer-associated fibroblast (CAF) or normal fibroblast (nlFB) annotation (top) and by D5-associated program score (bottom). **E)** Percentage of D5 program-positive/*NES/SOX4/NOTCH1*-positive fibroblasts per sample across normal, adjacent, and tumor tissues. Double-positive cells were defined as fibroblasts exceeding the 75th-percentile threshold for both the D5 program score and the *NES/SOX4/NOTCH1* signature score. Pairwise comparisons between tissue states were performed using Wilcoxon tests with Benjamini–Hochberg correction. **F)** Percentage of D5 program-positive/*NES/SOX4/NOTCH1*-positive fibroblasts per sample across normal, adjacent, and tumor tissues, shown separately for colorectal, gastric, lung, and PDAC samples. Pairwise comparisons between tissue states were performed within each cancer type using Wilcoxon tests with Benjamini–Hochberg correction. Comparisons for which either group contained fewer than three samples are indicated by the corresponding sample size rather than a P value. For **E** and **F**, samples containing fewer than 20 fibroblasts were excluded. Each point represents one sample.

We next examined whether the abundance of this double-positive population changed with tissue transformation. The frequency of late program-positive + *NES/SOX4/NOTCH1*-positive fibroblasts progressively increased from normal to adjacent to tumor tissue, with the highest abundance observed in tumors (all P < 0.001; **Fig. 5E**). To determine whether this pattern was restricted to a particular cancer type or shared across malignancies, we compared its frequency across normal, adjacent, and tumor tissue in organs represented in all three tissue states: colorectal, gastric, lung, and PDAC. In each organ, double-positive fibroblasts were nearly absent from normal tissue (0–0.12%), increased in adjacent tissue, and reached their highest frequencies in tumors (10.9–19.8%; **Fig. 5F**). This normal-to-adjacent-to-tumor gradient was observed consistently across all four organs and reached statistical significance in the majority of pairwise comparisons. Together, these findings show that the D5-associated transcriptional program is preferentially associated with CAFs and progressively enriched from normal to tumor tissue across multiple cancer types.

## DISCUSSION

Cancer-associated fibroblasts have been increasingly understood as dynamic populations whose activity is shaped by signals within the local tumor microenvironment rather than by fixed and irreversible subtype identities. In this study, we identify a reversible tumor-associated fibroblast program identifiable by expression of *NES, SOX4* and *NOTCH1* and define key signals and regulatory mechanisms controlling its activation. Our findings support a model in which this phenotype represents an environmentally maintained fibroblast state rather than a stable lineage. More broadly, it reinforces the idea that understanding CAF biology may require focusing less on defining discrete subtypes and more on identifying the signals that activate specific functional and transcriptional programs.

We identified EGF and FGF2 as sufficient drivers of the NES+SOX4+NOTCH1+ program in both patient-derived CAFs and human fibroblast cell lines. Importantly, these factors emerged from predicted tumor-to-fibroblast interactions in spatially resolved tissue, linking the *in vitro* response that we observed to signals present during disease progression. This is consistent with previous work showing that FGF2 can induce NESTIN expression in primary human dermal fibroblasts, supporting the broader ability of FGF signaling to activate neural-associated features in fibroblasts (32). Pancreatic tumor cells similarly induced NESTIN expression in nearby fibroblasts in the absence of exogenous growth factors, with the response decreasing with increasing distance from the tumor interface. These findings support a locally induced program driven by signals originating from the epithelial compartment. Furthermore, these findings are consistent with the concept that CAF states can be shaped by specific microenvironmental signals rather than representing fixed identities (14). The convergence of these pathways is particularly interesting considering recent work showing that EGFR and FGFR2 cooperate during pancreatic tumor initiation where dual inhibition reduces MAPK signaling, epithelial proliferation, and precursor-lesion formation (33). Interestingly, that study also showed that surrounding fibroblasts provide ligands required for epithelial FGFR2 activation, highlighting the reciprocal nature of growth-factor signaling between epithelial and stromal compartments. Our findings extend this concept in the opposite direction, showing that PDAC cells can in turn induce a growth factor–dependent transcriptional state in fibroblasts. Together, these observations suggest that EGF/FGF2 signaling may form part of a bidirectional tumor–stroma signaling network that contributes to pancreatic disease progression. The use of overlapping EGFR/FGFR signaling across epithelial and stromal compartments also raises the possibility that pathway redundancy may extend across cell types within the tumor microenvironment. Defining how these pathways are activated and compensated for in each compartment could therefore be important for understanding resistance to pathway-targeted therapies and for identifying more effective combination strategies.

A key feature of the induced state was its reversibility. Following growth factor withdrawal, NESTIN and TUJ1 expression returned toward baseline and the broader transcriptional profile shifted back toward that of untreated fibroblasts. This suggests that maintenance of the program depends on sustained extracellular signaling rather than stable lineage conversion. MAPK/ERK signaling provides a mechanistic link between growth-factor exposure and this reversible state. EGF/FGF2 stimulation altered phosphorylation of EGFR, RAF1, MEK1, ERK2 and downstream RSK proteins, while direct ERK inhibition produced the strongest shift away from the induced transcriptional program and toward the baseline fibroblast state. This is consistent with previous evidence that FGF2 regulates NESTIN through the Ras–Raf–ERK pathway in other cellular contexts (34) as well as with recent work showing that MAPK-high and MAPK-low malignant cell states in PDAC can rapidly interconvert following pathway inhibition (35).

The transcriptional response itself was also dynamic. Rather than moving directly from one stable state to another, fibroblasts progressed continuously through several phases, beginning with the loss of fibroblast- and ECM-associated features, followed by transient activation of a program enriched for neural-associated cytoskeletal genes and, later, a stress- and growth-arrest-associated program. This temporal organization is important when interpreting the individual markers used to define the phenotype. NESTIN, SOX4, NOTCH1 and TUJ1 did not respond identically to growth-factor withdrawal or pathway inhibition, suggesting that they are not interchangeable outputs of a single regulatory event. Instead, they appear to occupy different positions in time throughout the response. Similarly, we saw that the late acting factors SOX4 and NOTCH1, while both markers used in the definition of the phenotype were variably important in maintaining it. *SOX4* knockdown for example had little effect on the transcriptional program, consistent with its position as a leaf node in the late program adjacency matrix. While not necessary for the program itself, it could still be functionally relevant, as previous work in PDAC identified a SOX4–MMP11 axis in myCAFs that promotes tumor-cell invasion (36). NOTCH1 was even more interesting as inhibition appeared to decrease the phenotype, without having major effects on the programs, though knockdown did appear to increase the initial fibroblast program suggesting it may play a role in keeping the normal fibroblast program suppressed.

*ETS*-family transcription factors emerged as important regulators of this trajectory. *ETV4*, *ETV1* and related factors were activated early following growth-factor exposure, while *ETS2* and *ETV1* were associated with the later program, and knockdown of these factors reduced the corresponding program scores. Several of these factors are established downstream effectors of MAPK/ERK signaling, consistent with the central role of this pathway in our phosphoproteomic and inhibitor analyses. Previous work also connects FGF signaling to ETS-dependent fibroblast activation. Bordignon et al. showed that FGF2 and TGF-β establish opposing fibroblast programs and identified *ETV1* as a critical determinant of this duality, mediating the FGF response while suppressing TGF-β-induced genes (37). Importantly, differences in *ETV1* expression were also observed across independent CAF strains and between fibroblast populations in human squamous cell carcinoma stroma, supporting the idea that this regulatory axis is associated with distinct fibroblast states *in vivo* (*37*). Together with our results, this suggests that ETS factors may translate extracellular EGF/FGF2 signaling into distinct fibroblast transcriptional programs.

The broader relevance of this program is supported by its enrichment across multiple human cancers. The late program was preferentially expressed in CAFs, and fibroblasts positive for both the late program and the *NES/SOX4/NOTCH1* signature increased from normal to tumor-adjacent and tumor tissue across several cancer types. These findings suggest that the state is not restricted to pancreatic fibroblasts, although whether the same upstream signals and regulatory mechanisms drive it across tumor types remains to be determined. Its enrichment for *CDKN1A, GDF15, TIMP1* and *SERPINE2* along with the relative elevated activity of secreted growth factors at late time-points is consistent with increasing cellular stress or growth arrest following prolonged growth-factor exposure. Senescence has been described in pancreatic CAF populations and can be associated with distinct secretory and immunoregulatory properties (38,39). The fact that the program is fully reversible on growth factor withdrawal though, suggests that this is at least not a classical senescent state.

An important next step will be to define the functional consequences of this fibroblast state within the tumor microenvironment. Given its association with disease progression (26), future studies should determine how this state influences tumor cells and other components of the microenvironment, including whether it contributes to the establishment of an immunosuppressive niche. The neural-associated features of the program also raise the possibility of functional interactions with peripheral nerves or other neural components of the tumor microenvironment, an intriguing possibility that remains unexplored. Overall, our findings define the NES+SOX4+NOTCH1+ program as a reversible, growth-factor-dependent fibroblast state regulated through MAPK/ERK signaling to ETS-family transcription factors. These results provide a mechanistic link between tumor-derived signals and CAF plasticity and support the view that fibroblast heterogeneity can arise through dynamic, environmentally regulated transcriptional states. Determining the functional consequences of this state within the tumor microenvironment will now be important for understanding its contribution to disease progression.

## METHODS

### Cell Lines and culture conditions

Human fibroblast cell lines BJ (ATCC, cat# CRL-2522; neonatal male foreskin), MRC5 (ATCC, cat# CCL-171; male fetal lung), and IMR90 (ATCC, cat# CCL-186; female fetal lung), and the pancreatic cancer cell line SU8686 (ATCC, cat# CRL-1837; female ductal carcinoma) were used in this study. Fibroblast cell lines were maintained at passages below 10, and SU8686 cells were maintained below passage 25. BJ, MRC5, and IMR90 cells were cultured in DMEM (Sigma-Aldrich, cat# D6429) supplemented with 10% fetal bovine serum (FBS; Gibco, Thermo Fisher Scientific, cat# 12483020). SU8686 cells were maintained in RPMI-1640 (LifeTech, cat# 11875119) supplemented with 5% FBS. All cells were maintained at 37°C and 5% CO₂. (40,41)

All cell lines were confirmed to be mycoplasma-free prior to use by PCR-based testing using a custom protocol based on previously described methods (40,41). Briefly, 100 µL of cell culture supernatant was heat-treated at 95°C for 5 min and centrifuged at 10,000 × g for 1 min. Subsequently, 1 µL of the clarified supernatant was used as template in a PCR reaction with mycoplasma-specific primers. Amplification products were resolved on a 1% TAE agarose gel containing 0.25× GelRed (**Supplementary Tables: 1**).

Human myofibroblastic cancer-associated fibroblasts (myCAFs; f3870), inflammatory CAFs (iCAFs; f3910), and antigen-presenting CAFs (apCAFs; p4156) were derived from human pancreatic ductal adenocarcinoma (PDAC) or intraductal papillary mucinous neoplasm (IPMN) patient tissue by explant culture (26). Specifically, f3870 (myCAF) and f4156 (apCAF) were derived from a PDAC sample, f3910 (iCAF) from IPMN with low-grade dysplasia. All CAF subtypes were cultured in DMEM supplemented with 10% FBS, 1% penicillin-streptomycin, 2.25 mM glucose, 1× GlutaMAX (Gibco, Thermo Fisher Scientific, cat# 35050061), 1× N-2 supplement (Gibco, Thermo Fisher Scientific, cat# 17502048), 1× B-27 supplement (Gibco, Thermo Fisher Scientific, cat# 17504044), and 5 µg/ml gentamicin at 37°C with 5% CO₂.

For generation of fluorescently labelled cell lines, EGFP coding sequences were cloned into the dCAS9-VP64 backbone (Addgene, cat# 61422), replacing the dCAS9-VP64 insert and placing EGFP under the EF1α promoter. Lentiviral particles were produced in HEK293T cells by calcium phosphate co-transfection. Cells were seeded at 4 × 10⁶ cells per 10-cm dish in 8 mL DMEM supplemented with 10% FBS (four dishes per viral batch). The following day, cells were transfected with pMD2.G (Addgene, cat# 12259), psPAX2 (Addgene, cat# 12260), and transfer vector at a 1:1.8:2.3 molar ratio (0.72, 1.30, and 1.64 pmol per dish, respectively). DNA was mixed with CaCl₂ and an equal volume of 2× HEPES-buffered saline (140 mM NaCl, 1.5 mM Na₂HPO₄, 50 mM HEPES, pH 7.05) and added dropwise to cells. Medium was replaced the following day, and viral supernatant was collected 48 h post-transfection, pooled, filtered, and concentrated overnight at 4°C using 4× PEG/NaCl lentivirus concentrator (40% w/v PEG-8000, 1.2 M NaCl) at a 3:1 (v/v) supernatant-to-concentrator ratio. Virus was pelleted at 3,000 × g for 1 h at 4°C and resuspended in 100 µL Opti-MEM. BJ, MRC5, and IMR90 fibroblasts were reverse-transduced by combining 1 × 10⁵ cells with 1 µL concentrated virus in the presence of 0.4 µL/mL polybrene (Sigma, cat# TR-1003-G). Medium was refreshed after 24 h. At 72 h post-transduction, live GFP-positive cells were isolated by fluorescence-activated cell sorting on a BD FACSAria IIIu at the IRIC Flow Cytometry Core Facility.

### NES+SOX4+NOTCH1 program activation and reversibility assays

Fibroblast cell lines (BJ, MRC5, and IMR90) were cultured in Neurobasal medium (Gibco, Thermo Fisher Scientific, cat# 12348017) supplemented with 1× GlutaMAX, 1× N-2, and 2× B-27 (hereafter referred to as N3 medium), together with 20 ng/mL human recombinant EGF (StemCell Technologies, cat# 78006.1), 20 ng/mL human recombinant FGF2 (StemCell Technologies, cat# 78003), 10 ng/mL human recombinant BDNF (StemCell Technologies, cat# 78005), and 10 ng/mL human/mouse recombinant NT-3 (StemCell Technologies, cat# 78074). This complete induction medium is hereafter referred to as N3EFNB medium. Cells were maintained in N3EFNB medium for 5 days, with complete medium replenished every other day. Where indicated, growth factors were withdrawn on day 5 by replacing the induction medium with standard culture medium (DMEM supplemented with 10% FBS), and cells were maintained for an additional 3 days prior to analysis.

### Co-culture Assays

To recapitulate the ratio of cancer-associated fibroblasts to tumor cells observed in the pancreatic tumor microenvironment, co-cultures were established at a 5:1 fibroblast-to-tumor cell ratio. Briefly, 2,500 GFP-expressing fibroblasts (BJ, MRC5, or IMR90) and 500 mCherry-expressing SU8686 cells were seeded per well in 96-well plates and cultured for 5 days in DMEM/F-12 with GlutaMAX (Gibco, Thermo Fisher Scientific, cat# 10565018) supplemented with 1× N-2 and 2× B-27. This adapted medium was used in place of Neurobasal medium to support SU8686 viability during co-culture. Control conditions included SU8686 cells cultured without fibroblasts (tumor growth control), fibroblasts cultured in the same medium without growth factors (negative control for marker expression), and fibroblasts cultured in growth factor-supplemented medium (positive control for marker expression). Live-cell fluorescence imaging was performed every 12 h over the 5-day culture period using a Cytation 5 imaging reader (BioTek) at 4× magnification, capturing GFP and mCherry channels. Each fibroblast cell line was assessed in one well per experiment across independent experiments performed on different days (BJ and MRC5, n = 4; IMR90, n = 3). Tumor cell proliferation and spatial fibroblast marker expression were quantified as described in *Image Acquisition and Analysis*.

### Growth Factor Treatments and Inhibitor Experiments

Unless otherwise stated, all growth factor and inhibitor experiments were performed in the human fibroblast cell lines BJ, MRC5, and IMR90. For NES⁺SOX4⁺NOTCH1⁺ program activation, 2,000 cells per well were seeded in 96-well plates in N3EFNB medium and cultured for 5 days, with complete medium replenished every other day, as described in *Cell Lines and Culture Conditions*.

To identify the minimal growth factor combination required for NES+SOX4+NOTCH1+ program activation, a leave-one-out (N−1) experiment was performed. Cells were cultured in N3EFNB medium (positive control), N3 medium (negative control), or N3EFNB medium lacking individual growth factors (−EGF, −FGF2, −BDNF, or −NT3). Pairwise omission conditions lacking either EGF and FGF2 (−EGF/−FGF2) or BDNF and NT3 (−BDNF/−NT3) were also included. Dose-response experiments were subsequently performed with EGF and FGF2, either individually or in combination, at concentrations of 1, 5, 10, 20, and 40 ng/mL. For combined treatments, EGF and FGF2 were added at equivalent concentrations. All treatments were performed in N3 medium for 5 days with complete medium replenished every other day.

To assess the role of NOTCH signaling in the NES+SOX4+NOTCH1+ program activation, cells (including myCAFs) were treated with the γ-secretase inhibitors DAPT (20 µM; N-[N-(3,5-difluorophenacetyl-L-alanyl)]-S-phenylglycine t-butyl ester; Adooq, cat# A10288-5) or Nirogacestat (10 µM; Ambeed, cat# A421613-10MG) concurrently with N3EFNB medium. Inhibitors were maintained throughout the 5-day induction period. Vehicle control wells received an equivalent volume of anhydrous DMSO (0.1% v/v final concentration; Sigma, cat# 276855-100ML) in N3EFNB medium. Cells cultured in N3EFNB medium or standard culture medium (DMEM supplemented with 10% FBS) served as positive and negative controls, respectively.

To determine whether NOTCH ligands alone were sufficient to activate the NES+SOX4+NOTCH1+ program, recombinant NOTCH ligands were immobilized on 96-well plates. Plates were coated overnight at 4°C with recombinant human DLL1 (Sino Biological, cat# 11635-H02H), DLL4 (RCD Systems, cat# AVI10185-050), JAG1 (Sino Biological, cat# 11648-H02H), or JAG2 (MedChemExpress, cat# HY-P79094) diluted in PBS at the indicated concentrations. The following day, 1,000 myCAF cells or 2,000 fibroblast cell line cells per well were seeded onto coated plates in N3 medium. Uncoated wells containing N3EFNB medium served as a positive control for NES+SOX4+NOTCH1+ program activation, whereas uncoated wells containing N3 medium served as a negative control. Cells were cultured for 5 days with complete medium replenished every other day.

Additional signaling pathways were selected for inhibition based on phosphoproteomic analysis of EGF/FGF2-treated fibroblasts (see *Phosphoproteomics*). Cells were treated with inhibitors targeting AKT (Triciribine, 1 µM; Toronto Research Chemicals, cat# T774245; or Pictilisib dimethanesulfonate, 1 µM; MedChemExpress, cat# HY-20180), MEK1/2 (U0126, 1 µM; Cayman Chemical, cat# 70970-1), or ERK1/2 (Ulixertinib, 1 µM; MedChemExpress, cat# HY-15816). Inhibitor concentrations were selected based on western blot validation of target inhibition (**Supplementary Fig.6B**). Inhibitors were added concurrently with N3EFNB medium and maintained throughout the 5-day culture period. Vehicle control wells received 0.1% DMSO (v/v) in N3EFNB medium, while N3EFNB medium and standard culture medium served as positive and negative controls, respectively.

### Knockdown assay by siRNA transfection

Fibroblast cell lines (BJ, MRC5, and IMR90) were seeded onto Matrigel-coated 6-well plates at 1 × 10⁵ cells per well and cultured overnight in DMEM supplemented with 10% FBS at 37°C and 5% CO₂. The following day, medium was replaced with N3 medium, and cells were reverse transfected with Silencer™ Select siRNAs (Thermo Fisher Scientific), either individually or in combination. Combinatorial knockdowns included *ETV1+ETS2, ETV4+ETV5*, and *ETS2+ETV1+ETV4+ETV5*. Two independent siRNAs were used per gene: *SOX4* (s13300, s13301; cat# 4427037), *NOTCH1* (s9633, s9634; cat# 4392420), *ETS2* (s4850, s4851; cat# 4427037), *ETV1* (s4853, s4854; cat# 4392420), *ETV4* (s4861, s4862; cat# 4427037), and *ETV5* (s4863, s4864; cat# 4392420). A scrambled/non-targeting siRNA (Silencer™ Select Negative Control No. 1, cat# 4390843) was used as a control. Transfections were performed using Lipofectamine RNAiMAX (Thermo Fisher Scientific, cat# 13778030). siRNAs and Lipofectamine RNAiMAX were diluted separately in Opti-MEM (Gibco, Thermo Fisher Scientific, cat# 31985062), combined at a 1:1 ratio, and incubated for 5 min at room temperature before addition to cells. Each siRNA species was used at 5 pmol per well, including in multi-gene knockdown conditions, with transfection volume and Lipofectamine RNAiMAX scaled proportionally to the total number of siRNAs used. Cells were incubated with transfection complexes for 5 h at 37°C, after which the medium was replaced with N3EFNB medium to initiate induction. Complete medium was replenished every other day, as described in *Cell Lines and Culture Conditions*. Cells were collected on day 5 for RNA extraction and sequencing, as described in *Sample Preparation and Sequencing*.

### Cell lysates and western blotting

Cells were washed once with PBS then scraped directly in ice cold lysis buffer and transferred into microcentrifuge tubes. Lysis buffer was composed of 25 mM Tris-HCl pH 7.5, 150 mM NaCl, 1% IGEPAL, 5mM MgCl_2_, and contained 20 μM leupeptin and aprotinin (0.15 U/mL), 1 mM PMSF, phosphatase inhibitor cocktail (Millipore Sigma, cat# P0044), and 1 mM sodium orthovanadate. Cells were lysed on ice for 5 min, centrifuged at 22,000 x g at 4°C for 5 min, and boiled for 5 min after the addition of 5X sample buffer (300 mM Tris-HCl pH 6.8, 10% SDS, 0.1% bromophenol blue, 25% glycerol, 150 mM β-mercaptoethanol).

For immunoblotting, lysates were resolved on SDS–PAGE and proteins were transferred onto PVDF membranes (Bio-Rad, cat# 1620177). Membranes were blocked for at least 30 min in TBST (10 mM Tris-HCl pH 8.0, 150 mM NaCl, 0.2% Tween-20) containing 5% milk (BioShop, cat# SKI400.1) and then incubated with the following primary antibodies overnight at 4°C: anti-Phospho-AKT (1:2000; Cell Signaling Technology, cat# 4060), anti-AKT (1:2000; Cell Signaling Technology, cat# 4691), anti-Phospho-ERK (1:2000; Millipore Sigma; cat# M8159), anti-ERK1/2 (1:2000; Cell Signaling Technology, cat# 4695), anti-Phospho-p90RSK (1:1000; Cell Signaling Technology, cat# 8753), anti-RSK1/2/3 (1:1000; Cell Signaling Technology, cat# 9355). Secondary antibodies were anti-mouse-HRP (1:10,000; Jackson ImmunoResearch Laboratories Inc, cat# 115-035-146) and anti-rabbit-HRP (1:10,000; Jackson ImmunoResearch Laboratories Inc, cat# 111-035-144). Chemiluminescence signal was detected using homemade ECL and autoradiography film (Labscientific Inc., XAR ALF 2025).

### Immunofluorescence (IF)

Cells were fixed with 4% paraformaldehyde (PFA; Electron Microscopy Sciences/Cedarlane, cat# 15710) in PBS at 37°C for 30 min, followed by two washes with PBS. Cells were permeabilized for 10 min at room temperature with 0.1% Tween-20 (BioShop, cat# TWN510.100) in PBS supplemented with 2% FBS. Directly conjugated primary antibodies were diluted in PBS containing 2% FBS and 0.1% Tween-20 and incubated with cells overnight at 4°C. Antibodies used were Alexa Fluor 594 anti-Nestin (clone 10C2, 1:2,000, 0.25 µg/mL; BioLegend, cat# 656804), Alexa Fluor 647 anti-TUJ1 (clone TUJ1, 1:2,000, 0.25 µg/mL; BioLegend, cat# 801209), and Alexa Fluor 647 anti-SOX4 (clone B-7, 1:50, 1 µg/mL; Santa Cruz Biotechnology, cat# sc-518016). Following three washes with PBS supplemented with 2% FBS, nuclei were counterstained with DAPI (1 µg/mL; Sigma-Aldrich, cat# D9542). Imaging was performed using a Cytation 5 imaging reader (BioTek) at 4× magnification, acquiring DAPI, Texas Red (Nestin), Cy5 (SOX4 or TUJ1), and phase-contrast channels. Phase-contrast images were used exclusively for cell segmentation, as described in *Image Acquisition and Analysis*.

For experiments requiring sequential staining with antibodies detected in the same fluorescence channel, fluorescence was chemically quenched between staining rounds. A fresh quenching solution containing 9% hydrogen peroxide (BioShop, cat# HYP001.500) and 50 mM NaOH (BioShop, cat# SHY500.1) in PBS was mixed in equal volume with PBS supplemented with 2% FBS already present in the wells, yielding final concentrations of 4.5% hydrogen peroxide and 25 mM NaOH. Cells were incubated in quenching solution overnight at 4°C, washed twice with PBS supplemented with 2% FBS, and subjected to the subsequent staining round using the protocol described above.

### Quantitative PCR (qPCR)

Total RNA was extracted from approximately 1 × 10⁵ MRC5 fibroblasts collected at days 0, 2, and 5 of N3EFNB treatment using the PureLink RNA Mini Kit (Thermo Fisher Scientific, cat# 12183025) according to the manufacturer’s instructions. Day 0 samples were collected from cells maintained in standard culture medium (DMEM + 10% FBS) and served as the calibrator condition. RT-qPCR was performed at the IRIC Genomics Platform. In brief, RNA was treated with TURBO DNase (Thermo Fisher Scientific, cat# AM1907) to remove genomic DNA contamination and quality was assessed prior to cDNA synthesis using a Bioanalyzer 2100 (Agilent). Complementary DNA was synthesised using the Maxima First Strand cDNA synthesis kit (Thermo Fisher Scientific, cat# K1641).

Gene expression was determined using assays designed with the Universal Probe Library from Roche (www.universalprobelibrary.com). For each qPCR assay, a standard curve was performed to ensure that the efficiency of the assay was between 90% and 110% (**Supplementary Tables: 2)**. Reactions were run in duplicate on a QuantStudio qPCR instrument (Thermo Fisher Scientific). Relative expression comparison (RQ = 2^−ΔΔCt) was calculated using the ExpressionSuite software (Thermo Fisher Scientific), using HPRT and GUSB as endogenous controls. Three biological replicates were performed per condition using MRC5 fibroblast cell lines.

### Phosphoproteomics

To map signaling pathways activated by growth factor treatment, MRC5 fibroblasts were selected for phosphoproteomic analysis based on their robust response to induction (see *Results*). Cells were seeded at 5 × 10⁶ cells per 150 cm² dish 24 h before treatment. On the day of the experiment, culture medium was replaced with N3 medium, and cells were incubated for 3 h at 37°C and 5% CO₂ to establish baseline signaling conditions. EGF and FGF2 were then added simultaneously at 20 ng/mL each, and cells were harvested at 0, 10, 30, and 120 min post-treatment. Experiments were performed independently on three different days (n = 3 experimental replicates). At each timepoint, medium was removed and cells were washed twice with ice-cold PBS. Cells were scraped in 2 mL ice-cold PBS supplemented with phosphatase inhibitor cocktail (Sigma-Aldrich, cat# P0044; 20 µL/mL), transferred to 50-mL conical tubes, and centrifuged at 300 × g for 5 min at 4°C. Supernatants were discarded, and cell pellets were snap-frozen in liquid nitrogen and stored at −80°C until further processing. All steps following medium removal were performed on ice to preserve protein phosphorylation.

Cell pellets were lysed in 9 M urea, 100 mM Tris-HCl (pH 8.2), 10 mM TCEP (Thermo Scientific, cat# 77720), 40 mM chloroacetamide (Sigma-Aldrich, cat# C0267), and 1× phosphatase inhibitor cocktail (Sigma-Aldrich, cat# P0044). Lysates were sonicated at 15 W using three 10-s bursts with cooling on ice, vortexed for 1 h at room temperature, and cleared by centrifugation at 20,000 × g for 15 min. Protein concentration was measured using a NanoDrop, and 250 µg protein per sample was diluted to 1 M urea with 100 mM Tris-HCl (pH 8.2) and digested overnight with 5 µg sequencing-grade trypsin at 37°C with shaking at 500 rpm. Digests were acidified with 1% TFA, cleared by centrifugation at 13,000 × g for 10 min, desalted using HLB cartridges, and enriched for phosphopeptides using TiO₂ micro-columns (GL Sciences, cat# GL5010-21307) according to the manufacturers’ protocols. Phosphopeptides were dried, resuspended in 4% formic acid, loaded onto Evotips (Evosep), and separated on a 15-cm Aurora column (IonOpticks; 75 µm i.d.) using the 20 samples-per-day Whisper Zoom method on an Evosep Eno system coupled to an Orbitrap Ascend Tribrid mass spectrometer (Thermo Fisher Scientific). Data were acquired in data-dependent acquisition mode. Full MS spectra were acquired at 120,000 resolution with a 50-ms injection time, followed by HCD MS/MS spectra of the most abundant multiply charged precursors within a 3-s cycle using 27% collision energy, 30,000 resolution, a 50% AGC target, and a 59-ms injection time.

Raw phosphopeptide data were processed in-house in R (v4.4.1). Phosphopeptides containing at least one phosphorylation site (n = 19,842) were collapsed to 13,488 unique phosphosites by summing intensities across peptide forms mapping to the same site. Intensity values of 0 were considered undetected and treated as missing values. Intensities were log₂-transformed and median-centred within each sample, and log₂ fold-changes relative to t = 0 were calculated independently for each biological replicate. Phosphosites detected in all three replicate pairs (n = 9,963) were evaluated using a best-timepoint approach. For each site, the timepoint showing the largest absolute fold-change was selected and tested against zero using a one-sample t-test. Sites were considered regulated when p < 0.05, |log₂ fold-change| > 1, and the direction of change was concordant in at least two of three replicates, yielding 583 regulated sites. Because the tested timepoint was selected from the observed data and no multiple-testing correction was applied at this stage, p-values were considered descriptive and used for exploratory prioritisation. Sites undetected in all replicates of one condition were classified as ON (detected only after stimulation) or OFF and analysed separately. Pathway enrichment was performed using the 564 genes containing either a significantly regulated phosphosite or an ON site (378 and 186 genes, respectively). KEGG pathway enrichment was performed using clusterProfiler (enrichKEGG; v4.12.6) with a hypergeometric test, Benjamini–Hochberg adjusted p < 0.05, and the whole genome as background. Significantly enriched pathways were visualized as a network, with phosphoproteins assigned to their most significantly enriched pathway. Schematic relationships between selected enriched pathways were manually included when supported by KEGG pathway maps; these relationships were not inferred from the phosphoproteomic enrichment analysis (42,43).

### Single-cell RNA Sequencing and Analysis

#### Cell Fixation

BJ, MRC5, and IMR90 fibroblasts were collected at days 0, 2, and 5 of NES+SOX4+NOTCH1+ program activation and pooled with AML5 and NPC reference populations at each timepoint, generating three independent libraries for single-cell RNA sequencing. At day 0, fibroblasts were maintained in standard culture medium (DMEM supplemented with 10% FBS) prior to collection, whereas day 2 and day 5 samples were collected during induction as described in *Cell Lines and Culture Conditions*. Prior to library preparation, cells were fixed using a DSP-methanol fixation protocol (Fluent BioSciences, FB0004708). Briefly, up to 1 × 10⁶ cells were resuspended in 200 µL Cell Suspension Buffer (Fluent BioSciences, cat# FB0002440) on ice. A fresh DSP stock was prepared by dissolving dithiobis(succinimidyl propionate) (DSP; Pierce, Thermo Fisher Scientific, cat# A35393) in anhydrous DMSO (Thermo Fisher Scientific, cat# D12345) at 50 mg/mL. Twenty microlitres of DSP stock were added to 800 µL ice-cold 100% methanol (Sigma-Aldrich, cat# MET302.1), and the resulting fixative was gradually added to the cell suspension with gentle swirling. Cells were incubated on ice for 30 min, after which DSP was quenched with 20.4 µL of 1 M Tris-HCl (pH 7.5). Fixed cells were stored at −20°C until further processing. For recovery, fixed cells were centrifuged at 500 × g for 5 min at 4°C, the methanol-containing supernatant was removed, and pellets were resuspended in freshly prepared buffer containing 3× saline sodium citrate, 1 µM DTT, 0.2 U/µL NxGen RNase inhibitor (Biosearch Technologies, cat# 30281-1), 1% BSA (Sigma-Aldrich, cat# A1595), and nuclease-free water. Cells were filtered through a 40-µm strainer, recounted, and diluted to 3,400–4,250 cells/µL in Cell Suspension Buffer.

#### Reference Cell Populations

Two additional cell populations were included as reference controls. AML5 cells, an acute myeloid leukaemia cell line (DSMZ, cat# ACC-247), were cultured in RPMI-1640 supplemented with 10% FBS and served as a transcriptionally distinct reference population for batch correction across timepoints. Neural progenitor cells (NPCs) were differentiated from SBAD3 iPSCs obtained from StemBancc using the STEMdiff SMADi Neural Induction Kit (STEMCELL Technologies, cat# 08581) and maintained in STEMdiff Neural Progenitor Medium (STEMCELL Technologies, cat# 05833) according to the manufacturer’s instructions. NPCs were included as an external transcriptional reference to assess the similarity of growth factor-induced fibroblasts to a bona fide neural progenitor state.

#### Library Preparation and Sequencing

Single-cell 3′ RNA-seq libraries were prepared from 17,000 fixed cells per sample using the PIPseq T10 3′ Single Cell RNA Kit v5.0 (Fluent BioSciences) according to the manufacturer’s fixed-cell protocol. Cell lysis, mRNA capture, reverse transcription, and whole-transcriptome amplification were performed within Particle-templated Instant Partitions (PIPs). Library quality was assessed by Qubit fluorometry and Agilent 2100 Bioanalyzer High Sensitivity DNA analysis prior to sequencing. Three libraries were generated, corresponding to D0 (DSP2348-1), D2 (DSP2348-2), and D5 (DSP2348-3). Sequencing was performed using a NextSeq 2000 (100-bp single-end reads) and a NovaSeq 6000 (100-bp paired-end reads; 1,500 M clusters), yielding 259,192,757, 390,644,622, and 369,666,025 clusters for D0, D2, and D5, respectively. Reads were demultiplexed allowing one mismatch, and sequencing adapters were removed prior to downstream analysis.

#### Computational Analysis

Raw reads were processed using PIPseeker (v3.1; Fluent BioSciences) and aligned to the human reference genome GRCh38 (2022.04 index) to generate filtered gene expression matrices. A Seurat object was generated for each timepoint using a minimum threshold of three cells per feature and 200 features per cell (44) (Seurat v5.4.0). Mitochondrial read content was calculated as the proportion of reads mapping to genes with the *MT-* prefix. Data were log-normalised using a scale factor of 10,000, and the top 2,000 highly variable features were identified using the variance-stabilising transformation method. Dimensionality reduction was performed by principal component analysis (PCA), using the top 10 principal components for graph-based clustering (resolution = 0.5) and UMAP visualisation. Seurat objects from the three timepoints were subsequently merged and re-clustered for joint analysis.

#### Donor Deconvolution

Because all five cell populations were pooled prior to library preparation, donor deconvolution was performed using Souporcell (45) (v2.1; k = 5 clusters). Souporcell was run on cell-barcode-tagged BAM files generated using the PIPseeker barcode command, with the GRCh38 primary assembly as reference. Doublets identified by Souporcell were excluded from downstream analyses, retaining singlets only.

Cell identities were assigned using a combination of transcriptional and genotype information. IMR90 cells were unambiguously identified by female-specific gene expression (XIST-positive and RPS4Y1-negative), while AML5 and NPC populations were identified based on marker gene expression and used as reference populations for batch integration and as biological benchmarks for neural progenitor identity. The two remaining male populations, corresponding to BJ and MRC5, were distinguished by SNP-based genotyping. Souporcell genotype profiles were extracted for each cluster at all three timepoints and matched across days using pairwise Pearson correlations of SNP allele frequencies at shared variants. Candidate discriminating SNPs with opposite homozygous genotypes between the two populations were identified using a minimum read depth of 10 and genotype quality score ≥ 20.

To experimentally confirm cluster identity, PCR primers were designed using 500-bp genomic regions spanning each candidate SNP (250 bp upstream and downstream) extracted from GRCh38 (**Supplementary Tables: 3**). BJ and MRC5 cell lysates were amplified using these primers, and PCR products were Sanger sequenced at the IRIC Genomics Core. Resulting sequences were aligned to the reference genome to confirm the cell-line identity assigned to each Souporcell cluster.

#### Pseudotime Inference and Trajectory Embedding

Pseudotime was inferred independently for BJ, MRC5, and IMR90 fibroblasts using Slingshot (46) (v2.12.0). For each cell line, PCA was repeated using only cells from that line and the previously identified variable features. Cells were assigned to a single lineage using experimental timepoints (D0, D2, and D5) as cluster labels, with the trajectory explicitly rooted at D0 and terminating at D5. Pseudotime values were extracted for each cell and quantile-normalised within each cell line by clipping values to the 5th–95th percentile range and rescaling them from 0 to 1, allowing trajectories to be compared across cell lines.

For visualisation in a reduced-dimensional space independent of pseudotime assignment, classical multidimensional scaling (MDS) was performed on the Euclidean distance matrix derived from the top 10 principal components for each cell line. Cells were coloured according to their normalised pseudotime values.

For trajectory visualisation, cells from each line were divided into 10 pseudotime quantile bins. This binning scheme was used consistently throughout subsequent trajectory-based analyses. For each gene, regulon, or ligand-activity score, the median and interquartile range (25th–75th percentile) were calculated within each bin. A smoothing spline (smooth.spline, spar = 0.2) was fitted to the bin means to generate a continuous trajectory, with the interquartile range displayed as a shaded ribbon. A stacked bar beneath each trajectory shows the proportion of cells from each experimental timepoint (D0, D2, and D5) represented within each bin, allowing pseudotime progression to be interpreted in relation to experimental time.

#### Gene expression program discovery by consensus integrative NMF (cNMF)

To identify gene expression programs associated with the transcriptional response to N3EFNB treatment, we performed consensus integrative non-negative matrix factorization (ciNMF) using LIGER (rliger v2.2.1) on the merged D0/D2/D5 dataset, treating each cell line (MRC5, BJ, IMR90) and reference population (AML5, NPC) as a separate dataset within the LIGER framework. Data were log-normalized (scale factor 10,000) and the top 3,000 highly variable genes were identified (Seurat) prior to factorization. Integrative NMF was performed across a range of factor numbers (k = 6–20; λ = 5). The optimal factor number was selected by jointly evaluating pooled relative reconstruction error and factor redundancy, quantified as the maximum and median absolute Spearman correlation between per-cell factor usage scores. Reconstruction error decreased substantially between k = 6 and k = 12–13, after which improvements were modest and non-monotonic. Both maximum and median factor redundancy reached a minimum at k = 13 and generally increased at higher values of k. Based on the balance between model fit and factor redundancy, k = 13 was selected for downstream analysis (**Supplementary Fig. 3G-H**)

Per-cell loadings for all 13 factors were incorporated into the Seurat object; factors whose activity varied significantly over the induction time-course were identified as described below. The top loading genes for each factor were extracted from the shared loading matrix (W) and used to assign biological identity to each program.

#### SCENIC regulon inference

Raw counts, restricted to genes with valid HGNC gene symbols and detected in more than 100 cells, were exported to loom format (SCopeLoomR v 0.13.0). Transcription factor regulons and per-cell regulon activity were inferred using pySCENIC (v0.12.1) with default AUCell scoring. Candidate regulons were pruned for direct cis-regulatory evidence using RcisTarget against the human (hg38, RefSeq r80) mc9nr cisTarget motif databases, comprising both 10kb up/downstream-of-TSS and 500bp-upstream/100bp-downstream-of-TSS gene-based motif rankings, with motif-to-transcription-factor annotations from the motifs-v9-nr.hgnc-m0.001-o0.0 table. The curated human transcription factor list distributed with pySCENIC was used to restrict candidate regulators.

#### Ligand-activity scoring

Ligand activity was estimated per cell using the NicheNetv2 (47) ligand-target prior models (https://zenodo.org/records/7074291). For each ligand in the NicheNet ligand-target matrix, candidate target genes were ranked by regulatory potential, and up to the top 100 genes with a positive score were retained as that ligand’s target gene set, ligands with fewer than 5 such genes were excluded from downstream scoring. Ligands supplied exogenously in the induction medium (FGF2, EGF, BDNF, NTF3, insulin, transferrin) and laminin subunits were retained irrespective of expression filtering. All other candidate ligands were retained only if both the ligand and at least one of its receptors (per the NicheNet ligand-receptor network) were expressed in at least 5 cells in each of the three fibroblast cell lines (BJ, MRC5 and IMR90). Per-cell ligand activity was then scored for each retained ligand’s target gene set using UCell (v2.8.0) applied to raw counts.

#### Time-association testing of gene expression programs

To identify factors whose activity varied significantly over the induction time-course, we tested the association between each factor’s per-cell loading (ciNMF loading, SCENIC regulon activity, or ligand-activity score) and timepoint (D0, D2, D5) independently within each cell line, using a k-nearest-neighbor mutual information (MI) estimator (k=3). Statistical significance was assessed by comparison to a null distribution generated from 500 permutations of timepoint labels, with false discovery rate correction (Benjamini-Hochberg) applied within each cell line. For SCENIC and ligand activity derived scores, factors were additionally required to be significantly time associated (FDR ≤ 0.01) in all three fibroblast cell lines (MRC5, BJ and IMR90) and) and not significantly time-associated (FDR > 0.1) in the AML5 or NPC reference populations, to isolate regulators specific to the shared fibroblast response to N3EFNB induction

#### Identification of temporally coordinated regulon and ligand-activity clusters

Regulons and ligand-activity scores passing the fibroblast-specific time-association filter described above were binned into the same 10 pseudotime bins per line used above, and pairwise Pearson correlations were calculated between factors based on their per-bin mean trajectories (minimum correlation threshold = 0.7). Factors were represented as nodes in a k-nearest-neighbor graph (k = 5) and clustered using the Leiden algorithm (resolution = 0.5) to group factors with similar temporal activation patterns

#### Regulator coverage analysis of shared temporal programs

To identify candidate transcriptional regulators of the shared temporal gene expression programs, we intersected the top 100 loading genes of each shared, crossline time-associated ciNMF program (ciNMF1, ciNMF4, ciNMF2, corresponding to the D0, D2, and D5 stages of the shared fibroblast response) with SCENIC-inferred regulons (motif-pruned). For each program, SCENIC regulon target rows were merged by transcription factor, and TFs were ranked by the number of program genes contained in their merged target set (ties broken by smaller regulon size, then alphabetically). The top 10 regulators per program (23 unique TFs across the union of all three lists) were retained for visualization and network analysis. For each program, we calculated total gene coverage per regulator, the marginal number of new genes contributed when regulators were added in decreasing-coverage order, and pairwise containment between regulator target sets (the fraction of one regulator’s covered genes also covered by another) flagging candidate subset relationships where ≥80% of one regulator’s covered genes were also covered by a regulator with equal or greater coverage. Candidate TF–TF regulatory links were defined where the gene encoding one selected TF appeared within another selected TF’s merged regulon.

#### Bulk RNA-sequencing

Reversibility, signaling pathway inhibitor, and siRNA knockdown experiments were performed as described above (see *Cell Culture and Conditions* section) with cells collected for RNA extraction and sequencing at day 0, day 5, and day 8 (reversibility).

Cells were washed once with 1X PBS to remove residual growth medium and detached using Accutase (StemCellTechnologies, cat#7929), with exposure time minimized to preserve RNA quality. Cell suspensions were counted, centrifuged (300 × g, 5 min), and the supernatant removed. Pellets were resuspended in 50 µL of 1X DNA/RNA Shield (DNA/RNA Shield™ 2X Concentrate, Zymo Research, cat# R1200-25; diluted with nuclease-free water), with pellet volume kept below ∼10% of the total resuspension volume to avoid excess viscosity; samples that remained viscous after resuspension were vortexed for up to 2 min to shear genomic DNA. Samples were shipped at room temperature to Plasmidsaurus (Eugene, OR) for RNA extraction, library preparation, and sequencing with their RNA sequencing option (https://plasmidsaurus.com/technical-documentation/rna).

#### Reversibility analysis

To determine whether the reversibility observed at the protein level extended to the broader transcriptional program, bulk RNA-sequencing was performed on fibroblast cell lines (BJ, MRC5, IMR90) under control (D0, n=3 per line), N3EFNB-induced (D5, n=3 per line), and washout (n=1 per line) conditions. Differentially expressed genes (log2FC > 1, FDR < 0.01; **Supplementary Tables: 5**) between induced and control samples (n = 2,297 genes) were used for multidimensional scaling (leading log2FC method) of voom-normalized expression values.

#### N-1 Growth Factor Analysis

To determine whether EGF and FGF2 were sufficient to recapitulate the transcriptional changes induced by the complete N3EFNB cocktail, bulk RNA-sequencing was performed on BJ, MRC5 and IMR90 fibroblasts cultured under control, N3EFNB, EGF/FGF2 (EF), and EGF/FGF2/BDNF (EFB) conditions. N3EFNB-treated and control samples included three biological replicates per cell line, whereas EF and EFB conditions included one sample per cell line. Multidimensional scaling was performed on TMM-normalized log2-CPM expression values using the same set of differentially expressed genes identified between N3EFNB-treated and control samples in the reversibility analysis (|log2FC| ≥ 1, FDR ≤ 0.01). Treatment conditions were visualized separately for each fibroblast cell line to assess their transcriptional similarity to the complete N3EFNB-induced state.

#### Signaling pathway inhibitors analysis

To determine how inhibition of intracellular signaling pathways affected the transcriptional programs associated with N3EFNB induction, bulk RNA-sequencing was performed on BJ, MRC5 and IMR90 fibroblasts cultured for 5 days in N3EFNB medium in the presence of AKT (Triciribine), PI3K (Pictilisib), MEK (U0126), ERK (Ulixertinib), or NOTCH (DAPT or nirogacestat) inhibitors. Activity of the D0-, D2-, and D5-associated ciNMF programs (ciNMF-1, ciNMF-4, and ciNMF-2, respectively) was quantified from bulk RNA-sequencing profiles using UCell scores derived from the corresponding program gene sets. For each sample, program scores were corrected by subtracting the matched same-run, same-cell-line N3EFNB + DMSO control. D0 and DMSO samples were retained as visualization references but were not included as fitted inhibitor effects.

Inhibitor effects were estimated using Bayesian mixed-effects models with inhibitor condition as a fixed effect and cell line as a random intercept. A Student-*t* likelihood with three degrees of freedom was used, with zero-centered Student-*t* priors (df = 3, scale = 0.10) for inhibitor effects and Student-*t* priors (df = 3, scale = 0.05) for the cell-line and residual standard deviations. A separate experimental-run term was not included because treatment availability was confounded with run; instead, within-run DMSO subtraction was used as the primary batch correction.

#### siRNA analysis

To determine the contribution of candidate transcriptional regulators to the time-associated N3EFNB programs, bulk RNA-sequencing was performed on BJ, MRC5 and IMR90 fibroblasts following individual or combinatorial siRNA knockdown of *ETV1, ETV4, ETV5, ETS2, SOX4,* and *NOTCH1*. Knockdown efficiency was assessed directly from bulk RNA-sequencing expression values, and activity of the D0-, D2-, and D5-associated ciNMF programs (ciNMF-1, ciNMF-4, and ciNMF-2, respectively) was quantified using UCell. For each sample, the UCell score was corrected by subtracting the mean score of the matched same-run, same-cell-line scrambled siRNA control. D0, scrambled, and Lipofectamine-only controls were used as reference conditions and were excluded from model fitting.

Independent effects of each targeted gene were estimated using three parallel additive Bayesian mixed-effects models, one for each ciNMF program. Models included *ETV1, ETV4, ETV5, ETS2, SOX4,* and *NOTCH1* knockdown as independent binary predictors and cell line as a random intercept. Combination knockdowns were represented by the additive contribution of their constituent gene coefficients, without combination-specific terms or interactions. Models were fitted using a Student-*t* likelihood with three degrees of freedom and zero-centered Student-*t* priors (df = 3, scale = 0.03) for gene effects, cell-line standard deviation, and residual standard deviation. Run was not separately modeled because treatment availability was largely confounded with experimental run; same-run scrambled-control subtraction was therefore used as the primary batch correction. Evidence for an effect was summarized as the posterior probability that the coefficient fell outside a ±0.01 UCell-unit region of practical equivalence.

### UCell Scoring and Public Dataset Analysis

Single-cell RNA-seq data were obtained from a published pan-cancer atlas (11) (GEO: GSE210347), comprising 855,271 cells from 234 samples across 11 tumor types and normal, adjacent, and tumor tissue states. Samples were merged, log-normalised using Seurat, and batch-corrected across samples using fastMNN (batchelor v1.24.0; d = 30), as previously described. Clustering and UMAP visualisation were performed on the corrected embedding using Seurat.

Gene signatures corresponding to the day 2 (ciNMF-4) and day 5 (ciNMF-2) induction programs were derived from the integrative NMF analysis described above (*k* = 13). For each factor, the top 100 genes were selected based on loading weights in the shared W matrix. Program activity was initially scored across the entire tumor microenvironment (TME) using UCell (v2.12.0) to identify the cell populations in which each program was most highly expressed. Fibroblasts were then subset for downstream analysis (n = 104,804 cells) and annotated as cancer-associated fibroblasts (CAFs) or normal fibroblasts (nlFBs) based on cluster identity. The day 5 program score was subsequently recalculated using UCell within the fibroblast subset. Separately, a three-gene neural marker signature comprising *NES, SOX4,* and *NOTCH1* was scored using UCell after excluding genes detected in <1% of cells within the population being analysed. Cells were classified as double-positive when both the day 5 program score and neural marker score exceeded the 75th percentile of their respective distributions across all fibroblasts.

Co-occurrence between day 5 program positivity and neural marker positivity was assessed using mixed-effects logistic regression implemented in lme4, with neural marker positivity modelled as a function of day 5 program positivity and with random intercepts and random slopes for sample to account for within-sample non-independence. As a complementary per-sample summary, odds ratios were calculated for samples containing at least 50 fibroblasts using a 0.5 continuity correction.

For comparisons across tissue states (normal, adjacent, and tumor), the proportion of double-positive fibroblasts was calculated for each sample. Samples containing fewer than 20 fibroblasts were excluded. When multiple samples from the same patient represented the same tissue state, percentages were averaged so that each patient contributed a single value per tissue state. Patient identity was inferred from sample names provided in the original studies. Tissue states were compared using two-sided Wilcoxon rank-sum tests with Benjamini–Hochberg correction within each analysis panel. In organ-level analyses, comparisons involving a group with fewer than three patients were reported as n.d.

### Re-analysis of Ligand–Receptor Interactions from Xenium Spatial Transcriptomics

Xenium spatial transcriptomics data generated from Sections of 4 TMAs of 8 formalin-fixed paraffin-embedded samples spanning normal tissue through invasive intraductal papillary mucinous neoplasm (IPMN)-associated pancreatic carcinogenesis were re-analysed to investigate ligand–receptor interactions between tumor cells and fibroblasts (26). Regions of interest had previously been annotated into six histological categories reflecting disease progression: normal acinar tissue (N), acinar tissue (Ac), acinar-to-ductal metaplasia (ADM), low-grade dysplasia (LG), high-grade dysplasia (HG), and invasive carcinoma (IN). Cell-type annotation and spatial topic modelling using SpaTopic (48) (v1.3.1; 14 topics; Gibbs sampling) were performed as described in the related work (26).

For the present analysis, topic 6 was selected as the niche of interest based on two criteria. First, its abundance was associated with tumor grade progression in a parallel spatial modelling analysis. Second, its cellular composition was dominated by myCAFs and tumor cells, particularly the Tumoral_Invasive_S100P_CD166_CEACAM6 population (26).

Ligand–receptor interaction scores within topic 6 were computed separately for each sample using the spatCellCellcom function in Giotto (49) (v4.0.2). This method tests whether ligand– receptor pair expression between spatially proximal cell types exceeds that expected under a spatially permuted null distribution. For each ligand–receptor pair, a permutation interaction (PI) score was calculated as the log₂ fold-change between observed and permuted expression multiplied by −log₁₀ of the FDR-adjusted p-value, providing a combined measure of interaction magnitude and statistical confidence.

Interactions were restricted to tumor–fibroblast pairs in both directions and combined across samples within each histological grade using Stouffer’s method. P-values were sign-directed according to the PI score, with a rank-based correction applied to values rounded to zero by the permutation procedure. Combined p-values were adjusted for multiple testing within each grade using the Benjamini–Hochberg procedure, and ligand–receptor pairs with FDR < 0.05 were considered significant. Histological grades represented by fewer than three samples were excluded from grade-level analyses (Normal, n = 1).

Among significant tumor–fibroblast interactions, a subset of 13 ligands with established roles in neural biology was selected for further analysis. Interaction trends for these ligands were examined across histological grades. Because this subset was selected after significance testing, no additional multiple-testing correction was applied within the subset; reported FDR values correspond to the grade-level correction described above.

### Cerebral Organoid Experiments

Human dorsal forebrain cerebral organoids were generated from SBAD3 iPSCs using the STEMdiff Dorsal Forebrain Organoid Differentiation Kit (STEMCELL Technologies, cat# 08620) and maintained beyond day 43 in STEMdiff Neural Organoid Maintenance Kit (STEMCELL Technologies, cat# 100-0120) according to the manufacturer’s instructions. Cell transplantation into cerebral organoids was performed as previously described (31). Briefly, BJ, MRC5, and IMR90 fibroblasts were exposed to neural induction medium for 3 days prior to injection, as described in *Cell Lines and Culture Conditions*. GFP-expressing fibroblasts (50,000 cells in 2 µL) were injected into each organoid using an insulin syringe under a dissecting microscope. Cells were resuspended in 3 mg/mL growth factor-reduced Matrigel (Corning, cat# CLS356231) to facilitate retention at the injection site. iPSC-derived GFP-expressing NPCs were injected at the same cell number as a positive control, while mock-injected organoids receiving Matrigel without cells served as negative controls. Injected organoids were assessed at 1 week and 1-month post-injection by live-cell fluorescence imaging using a Cytation 5 imaging reader (BioTek) at 4× magnification in the GFP channel.

At the experimental endpoint, organoids were fixed in 10% formalin for 24 h at room temperature, paraffin-embedded, and sectioned at 15 µm using a microtome. Sections were mounted on glass slides and dried overnight at 42°C. Slides were deparaffinised through graded toluene and ethanol washes, rehydrated, and subjected to heat-mediated antigen retrieval in citrate buffer (10 mM citric acid, 0.05% Tween-20, pH 6.0) at 95–100°C for 30 min. Sections were then subjected to cyclic immunofluorescence, as described in *Immunofluorescence*, using two sequential staining rounds with DAPI nuclear counterstaining. Round 1 included anti-GFP (clone FM264G, 1:200, 2.5 µg/mL; BioLegend, cat# 338008), Alexa Fluor 594 anti-Nestin (clone 10C2, 1:2,000, 0.25 µg/mL; BioLegend, cat# 656804), and Alexa Fluor 647 anti-TUJ1 (clone TUJ1, 1:2,000, 0.25 µg/mL; BioLegend, cat# 801209). Following fluorophore quenching using the H₂O₂/NaOH protocol described in *Immunofluorescence*, round 2 included anti-MAP2 (clone SMI52, 1:1000, 0.5 µg/mL; BioLegend, cat# 801804) and anti-GFAP (clone SMI 25, 1:500, 1 µg/mL; BioLegend, cat# 837510). Images from sequential staining rounds were registered in FIJI using the Register Virtual Stack Slices plugin with a rigid transformation model.

#### Image Acquisition and Registration

All fluorescence imaging of cell culture experiments was performed using a Cytation 5 imaging reader (BioTek) at 4× magnification. For immunofluorescence experiments, images were acquired in the DAPI, Texas Red (NESTIN), Cy5 (SOX4 or TUJ1), and phase-contrast channels. Multi-round immunofluorescence experiments were imaged sequentially as described in *Immunofluorescence*. For multi-round staining experiments, cross-round image registration was performed using DAPI as the reference channel with custom scripts (https://github.com/djhfknapp/CycIF_ImageRegistration). Briefly, key points were detected in paired images using the Scale-Invariant Feature Transform (SIFT) algorithm and matched by brute-force descriptor comparison. An affine transformation was then estimated using the RANSAC algorithm and applied to all channels from the subsequent staining round to achieve spatial alignment across rounds. Marker identity was retained through channel metadata.

#### Monoculture Immunoffuorescence Analysis

A custom image-analysis pipeline was developed using FIJI (v1.54), Ilastik (v1.4.0), CellProfiler (v4.2.6), Python, and R (v4.4.1). Raw images first underwent illumination correction by rolling-ball background subtraction (radius = 50 pixels) in FIJI. The DAPI channel was independently brightness-normalised to improve nuclear segmentation across wells and imaging sessions. Cell bodies were segmented by pixel classification in Ilastik using phase-contrast images, generating per-pixel probability maps that were thresholded to produce binary segmentation masks. Nuclear segmentation was performed in CellProfiler using the brightness-adjusted DAPI channel to define primary objects, with cell bodies defined as secondary objects surrounding the segmented nuclei. Mean fluorescence intensity for each marker channel was extracted on a per-cell basis. Cells were classified as marker-positive using a threshold defined as the midpoint between the 95th percentile of the negative-control population and the 50th percentile of the positive-control population, calculated separately for each cell line and experiment. Results are reported as the percentage of marker-positive cells per well.

#### Co-culture Distance Analysis

For co-culture experiments, fibroblast and tumor cell populations were segmented independently from their respective fluorescence channels using population-specific Ilastik pixel-classification models. Per-cell measurements, including centroid coordinates, object area, and mean fluorescence intensity, were extracted in CellProfiler.

Spatial analysis of fibroblast marker expression relative to tumor cell proximity was performed using a custom Python script. For each fibroblast, the distance to the nearest tumor cell edge was calculated as the centre-to-centre distance minus the estimated radius of the nearest tumor cell. Tumor-cell radius was approximated from measured object area, and nearest neighbours were identified using a k-d tree spatial index implemented with scipy.spatial.cKDTree. Distances were converted from pixels to micrometres using a calibrated pixel size of 1.744 µm/pixel at 4× magnification. Fibroblasts were grouped into five proximity bins according to distance from the nearest tumor cell edge: 0–50, 50–100, 100– 150, 150–200, and 200–300 µm. Fibroblasts located more than 300 µm from the nearest tumor cell were excluded from distance-stratified analyses. Marker-positivity thresholds were derived from within-plate controls as described in *Monoculture Immunofluorescence Analysis*, and the percentage of marker-positive fibroblasts was calculated for each distance bin within each well.

#### Cerebral Organoid Image Acquisition and Analysis

For live-cell imaging of cerebral organoids, whole-organoid fluorescence images were acquired using a Cytation 5 imaging reader (BioTek) at 4× magnification in the GFP channel at designated timepoints. Mock-injected organoids were imaged first to establish exposure settings and minimise autofluorescence. Rolling-ball background subtraction (radius = 50 pixels) was applied in FIJI prior to image display to correct for variable background intensity across organoids.

Histological sections of paraffin-embedded organoids were imaged using Gen5 software (BioTek). Z-stacks were acquired at 4.2-µm intervals through the full 15-µm tissue section and combined using the default focus-stacking parameters. Multi-round immunofluorescence images were registered in FIJI using the Register Virtual Stack Slices plugin with a rigid transformation model (translation and rotation), using DAPI as the reference channel across staining rounds, as previously described (31). Marker-positive cells in histological sections were identified by manual review of registered multi-round immunofluorescence images. Display ranges were kept constant between injected and mock-injected control organoids to enable direct visual comparison.

### Bayesian Mixed-Effects Model

Bayesian mixed-effects models were used to quantify the effect of growth factor omission and signaling inhibition on marker expression across multiple cell lines and biological replicates simultaneously, using the brms package (v2.23.0) in R (v4.4.1) with Stan as the backend sampler. Prior to Bayesian fitting, pairwise interaction terms were evaluated by BIC comparison using lme4 (v1.1.38) and lmerTest (v3.2.1) and retained if ΔBIC < −2 relative to the additive base model. All models included cell line and biological replicate as random intercepts and were fitted with 4 chains of 4,000 iterations (1,000 warmup), adapt_delta = 0.99, and maximum tree depth = 15, with convergence assessed by R-hat and effective sample sizes.

For the N-1 growth factor omission experiment (NESTIN, SOX4, and TUJ1), the percentage of marker-positive cells was modeled directly using a Gaussian family with identity link. Weakly informative priors were specified: Normal (0, 20) for fixed-effect coefficients, Normal (0, 30) for the intercept, and half-Cauchy (0, 10) for standard deviation parameters.

For the signaling inhibitor experiments (NESTIN, SOX4, NOTCH1, and TUJ1), posterior predictive checks under the Gaussian model revealed a bimodal observed distribution not reproduced by the model, consistent with a subpopulation of cells that did not respond to induction. Marker expression was therefore modeled using a zero-inflated Beta family with logit link, which jointly estimates the expected marker expression among responding cells and the probability of non-response. Raw percentages were rescaled to proportions on [0, 0.9999) to satisfy the support of the Beta distribution. Weakly informative priors on the logit scale were specified: Normal (0, 2) for fixed-effect coefficients and intercepts, and Cauchy- (0, 2) for standard deviation parameters. We report fixed-effect coefficients for the response magnitude component, reflecting the effect of each condition on marker expression among responding cells.

For each model, fixed-effect coefficients were summarised as posterior mean, median, and 95% credible interval. An effect was considered directional if the posterior probability exceeded 0.9 in either direction. Results were visualised as forest plots of posterior density distributions with shaded 95% credible intervals.

### Statistical Analysis

All statistical analyses were performed in R (v4.4.1). For experiments comparing multiple treatment conditions against a single reference, a linear mixed-effects model was fitted using lme4 (v1.1.38) with biological replicate as a random intercept, followed by Dunnett’s multiple comparison test using the multcomp package (v1.4.29); when multiple cell lines were analysed jointly, cell line was additionally included as a random intercept. This approach was applied to NESTIN, SOX4 and TUJ1 expression in the fibroblast neural program activation experiment, NESTIN expression in the NOTCH inhibitor and NOTCH ligand experiments; and NESTIN and TUJ1 expression in the NES+SOX4+NOTH1+ program plasticity, and dose-response experiments. Unless otherwise stated, box plots show the median and interquartile range, with whiskers extending to 1.5 × the interquartile range.

For qPCR validation of scRNA-seq-derived temporal gene expression (NES, TUJ1, SOX9, SOX6, SOX4, ACTA2) during NES+SOX4+NOTH1+ program induction, expression changes at each timepoint (D2 and D5, relative to D0) were assessed by one-sample t-tests against zero, with Benjamini-Hochberg correction applied across genes.

CAF lines (myCAF, iCAF, apCAF) were analyzed separately, as each displayed different behaviour. For two-condition comparisons within individual CAF lines, Wilcoxon signed-rank tests were applied per line with Benjamini-Hochberg false discovery rate correction across lines.

For the co-culture experiments, the distance-dependence of NESTIN expression was assessed in two ways. First, for each well, a linear regression of the percentage of NESTIN+ fibroblasts against distance from the tumor interface (bin midpoints) was fitted. Second, the difference in the percentage of NESTIN+ fibroblasts between the zone adjacent to the tumor interface (0–50 µm) and the most distal zone (200–300 µm) was calculated. Slopes and paired differences were each analysed with an intercept-only linear mixed-effects model with experiment as a random intercept, to account for wells from the same experiment sharing culture conditions (lmerTest v3.2.1, Satterthwaite degrees of freedom; 95% confidence intervals from the Satterthwaite t distribution). Wells from all three fibroblast cell lines were analysed together.

Statistical significance was assessed at p < 0.05. Exact p-values are reported on figures, for comparisons with p-values between 0.001 and 0.05; comparisons with p < 0.001 are indicated as such. Non-significant comparisons are labelled “ns”.

### Ethics approval for use of human samples

Ethics approval for the use of relevant human cells and samples for this study was granted by the Comité d’éthique de la recherche clinique (CERC) of the Université de Montréal and the Comité d’éthique à la recherche (CÉR) at the Centre de recherche du CHUM (CRCHUM).

## Data availability

All raw data will be made available upon peer-reviewed publication.

## ACKNOWLEDGEMENTS

We thank Éric Bonneil and Mirela Pascariu from the Proteomics Core Facility at the Institute for Research in Immunology and Cancer (IRIC) for processing the phosphoproteomics samples; Raphaëlle Lambert from the Genomics Core at IRIC for sequencing; Mélina Narlis from the Cancer Research Histology Core at IRIC; and Annie Gosselin and Angélique Bellemare-Pelletier from the Flow Cytometry Core at IRIC for their technical support.

Funds for this work were provided through the IRIC Philanthropic funds from the Marcelle and Jean Coutu foundation and from the Fonds de recherche du Québec - Santé (FRQS #295647). D.J.H.F.K. has salary support from FRQS in the form of a Chercheurs-boursiers Junior 1 fellowship (#283502). M.I.I.R. was supported by a Doctoral Research Award from Cancer Research Society in in partnership with the Canadian Institutes od Health Research-Institute of Cancer Research, an IRIC Doctoral Award from the Institut of Research in Immunology and Cancer, a Bourse de passage accélère de la maitrise au doctorat from the University of Montreal, and Bourse de Mérite aux cycles supérieurs.

## AUTHOR CONTRIBUTIONS

Myriam Iliana Ibanez Rios: Methodology, Software, Validation, Formal analysis, Investigation, Data Curation, Writing - Original Draft, Writing - Review & Editing, Visualization; Maëlle Batardiere: Software, Investigation, Formal analysis, Writing - Review & Editing; Elham Dianati: Investigation, Data Curation, Writing - Review & Editing; Guillaume Serwe: Investigation, Data Curation, Writing - Review C Editing; Sajanth Kanagasingam: Investigation, Data Curation, Writing - Review & Editing; Alexandre Archambault-Marsan: Investigation, Data Curation, Writing - Review & Editing; Marc Terrien: Supervision, writing – review & editing; Pierre Thibault: Supervision, writing – review C editing; Vincent Quoc-Huy Trinh: Validation, Supervision, Resources, Data Curation, Writing - Review & Editing, Funding acquisition; David J.H.F. Knapp: Conceptualization, Methodology, Software, Validation, Formal analysis, Resources, Data Curation, Writing - Original Draft, Writing - Review & Editing, Visualization, Supervision, Funding acquisition.

## SUPPLEMENTARY FIGURE LEGENDS

Figure S1. **Tumor-to-fibroblast signaling increases across IPMN progression and N3EFNB differentially induces CAF subtypes. A)** Tumor-to-fibroblast ligand–receptor interaction networks within the grade-associated topic 6 at acinar metaplasia (ADM), low-grade and high-grade stages. Nodes represent significant ligands, colored by mean PI score; edges connect tumor-derived ligands to fibroblasts. **B)** Representative immunofluorescence images of patient-derived apCAFs and iCAFs after 5 days in control or N3EFNB medium, stained for NESTIN (magenta), SOX4 (green) and DAPI (blue). iCAFs did not survive N3EFNB treatment and were not quantified. Scale bar, 50 µm.

Figure S2. **EGF and FGF2 drive a reversible NES+SOX4+NOTCH1+ program in fibroblasts. A)** Representative immunofluorescence images of BJ and IMR90 fibroblasts after 5 days in control or N3EFNB medium, stained for NESTIN (magenta), SOX4 (green), NOTCH1 (cyan), TUJ1 (yellow) and DAPI (blue). Scale bar, 50 µm. **B)** Representative immunofluorescence images of MRC5 fibroblasts after 5 days in control or N3EFNB medium, stained for TUJ1 (yellow) and DAPI (blue). Scale bar, 50 µm. **C)** Percentage of TUJ1+ BJ, MRC5 and IMR90 fibroblasts after 5 days in control or N3EFNB medium. Marker positivity was defined relative to within-plate negative controls (see Methods). Each dot represents one experimental replicate (n = 8 per cell line). Linear mixed-effects model with experimental replicate and cell line as random intercepts, followed by Dunnett’s test versus control. **D)** Representative immunofluorescence images of MRC5 and IMR90 fibroblasts in 5-day co-culture with SU8686 tumor cells, stained for NESTIN (yellow) and DAPI (blue). Scale bar, 50 µm. **E)** Representative immunofluorescence images of BJ and IMR90 fibroblasts in control medium, after 5 days of N3EFNB treatment, and after 5 days in N3EFNB medium followed by 3 days in control medium (DMEM+10%FBS; washout). Stained for NESTIN (magenta), TUJ1 (yellow) and DAPI (blue). Scale bar, 50 µm. **F)** Representative immunofluorescence images of BJ and IMR90 fibroblasts cultured for 5 days in control, N3, N3EFNB, or N3EFNB lacking NT3, BDNF, EGF, FGF2, EGF/FGF2 or NT3/BDNF, stained for NESTIN (magenta), SOX4 (green), TUJ1 (yellow) and DAPI (blue). Scale bar, 50 µm. **G)** Representative immunofluorescence images of MRC5 fibroblasts under the same growth-factor omission conditions as in (**F**), stained for TUJ1 (yellow) and DAPI (blue). Scale bar, 50 µm. **H**) Percentage of TUJ1+ fibroblasts following growth-factor omission. Forest plots show the posterior distribution of each effect relative to N3EFNB from Bayesian mixed-effects models with biological replicate and cell line as random intercepts. Dark shading indicates the 95% credible interval, and vertical lines indicate posterior medians (BJ and MRC5, n = 4; IMR90, n = 3 biological replicates). **I)** Percentage of NESTIN+ BJ, MRC5 and IMR90 fibroblasts after 5 days of treatment with EGF, FGF2 or EGF/FGF2 at the indicated concentrations (n = 3 biological replicates per cell line). Linear mixed-effects model with biological replicate and cell line as random intercepts, followed by Dunnett’s test versus control.

Figure S3. **Validation of single-cell identities, pseudotime trajectories and ciNMF model selection. A)** UMAP of the single-cell RNA-sequencing dataset colored by experimental timepoint, sex, Souporcell cluster and assigned cell type. **B)** UMAP feature plots showing expression of *RPS4Y1, XIST, SOX2, PAX6, PTPRC* and *SPI1* used to support cell-type and sex assignments. **C)** Genotype-correlation matrices between Souporcell clusters across D0, D2 and D5, showing concordance of inferred genotypes across timepoints. **D)** Representative Sanger sequencing traces from BJ and MRC5 cells used to validate genotype-based cell-line assignments. **E)** UMAP of cells colored by final cell-type annotation. **F)** MDS embeddings of MRC5, BJ and IMR90 fibroblasts, from left to right, colored by normalized pseudotime. **G)** Pooled relative reconstruction error across ciNMF solutions with k = 6–20. **H)** Maximum and median factor redundancy across ciNMF solutions with k = 6–20. k = 13 was selected for downstream analysis based on the balance between reconstruction error and factor redundancy.

Figure S4. N3EFNB-treated fibroblasts do not acquire a bona fide neural progenitor phenotype and exhibit additional pseudotime-associated regulatory patterns. A) Representative GFP fluorescence images of cerebral organoids following vehicle injection or transplantation with GFP-expressing MRC5 fibroblasts or iPSC-derived NPCs, assessed at 1 and 4 weeks after injection. MRC5 fibroblasts were exposed to N3EFNB for 3 days before transplantation. Organoid outlines are shown in black. Scale bar, 100 µm. **B)** Representative immunofluorescence images of a cerebral organoid 1 week after transplantation with GFP- expressing MRC5 fibroblasts, showing regions proximal and distal to the injection site. Sections were stained for GFP, NESTIN, TUJ1, GFAP and MAP2. GFP+ regions are outlined in faint white lines. **C)** Pseudotime-resolved transcription factor regulon activities in MRC5, BJ and IMR90 fibroblasts, grouped into suppressed, transiently active and late-activated temporal patterns during N3EFNB treatment. **D)** Pseudotime-resolved activities of regulons and inferred ligand activities not assigned to the major temporal clusters shown in **Figure 3E** (“orphan activities”) across MRC5, BJ and IMR90 fibroblasts.

Figure S5. **Time-associated gene programs and regulatory activities during N3EFNB treatment. A)** Heatmaps of the top genes associated with the D0-, D2- and D5-associated ciNMF programs (ciNMF-1, ciNMF-4 and ciNMF-2, respectively) across MRC5, BJ and IMR90 fibroblasts at D0, D2 and D5 of N3EFNB treatment. Values represent row-scaled z-scores. **B)** Heatmaps of the top time-associated transcription factor regulon activities across MRC5, BJ and IMR90 fibroblasts at D0, D2 and D5. Values represent row-scaled z-scores. **C)** Heatmaps of the top time-associated inferred ligand activities across MRC5, BJ and IMR90 fibroblasts at D0, D2 and D5. Values represent row-scaled z-scores.

Figure S6. **Pathway inhibition differentially affects signaling and NES+SOX4+NOTCH1+ marker expression during N3EFNB induction. A)** Representative immunofluorescence images of BJ and IMR90 fibroblasts cultured for 5 days under the indicated conditions and stained for NESTIN (magenta), SOX4 (green), NOTCH1 (cyan), TUJ1 (yellow) and DAPI (blue). Scale bar, 50 µm. **B)** Western blot analysis of phosphorylated and total ERK1/2, AKT and RSK1/2/3 in BJ, MRC5 and IMR90 fibroblasts cultured under control or N3EF conditions with DMSO, Triciribine, Pictilisib, Ulixertinib or U0126. **B)** Representative immunofluorescence images of MRC5 fibroblasts cultured for 5 days under the indicated conditions and stained for TUJ1 (yellow) and DAPI (blue). Scale bar, 50 µm. **D)** Percentage of TUJ1+ BJ, MRC5 and IMR90 fibroblasts following 5 days of treatment with pathway inhibitors. Marker positivity was defined relative to within-plate negative and positive controls (see Methods). Bayesian mixed-effects models included biological replicate and cell line as random intercepts. Dark shading indicates the 95% credible interval, and vertical lines indicate posterior medians (BJ, n = 3; MRC5 and IMR90, n = 4 experimental replicates).

Figure S7. **NOTCH signaling contributes to NESTIN induction but is not sufficient to activate the NES+SOX4+NOTCH1+ program. A)** Representative immunofluorescence images of BJ, MRC5 and IMR90 fibroblasts cultured for 5 days in N3EFNB medium with vehicle, DAPT (20 µM) or nirogacestat (10 µM), stained for NESTIN (magenta) and DAPI (blue). Scale bar, 50 µm. **B)** Percentage of NESTIN+ fibroblasts from **(A).** Marker positivity was defined relative to within-plate negative and positive controls (see Methods). BJ and IMR90, n = 4; MRC5, n = 4–5 biological replicates. Linear mixed-effects model with biological replicate and cell line as random intercepts, followed by Dunnett’s test versus N3EFNB + vehicle. **C)** Percentage of NESTIN+ BJ, MRC5 and IMR90 fibroblasts cultured for 5 days in N3 medium (Neurobasal medium with N-2 and B-27 supplements) on plate-bound recombinant DLL1 (15 µg/mL), DLL4 (15 µg/mL), JAG1 (15 µg/mL) or JAG2 (5 µg/mL), or in uncoated wells with N3 or N3EFNB as negative and positive controls, respectively. n = 3 biological replicates per cell line. Linear mixed-effects model with biological replicate and cell line as random intercepts, followed by Dunnett’s test versus N3 control. **D)** Activity of the D0-, D2- and D5-associated ciNMF programs in bulk RNA-sequencing samples from BJ, MRC5 and IMR90 fibroblasts cultured for 5 days in N3EFNB medium with DAPT or nirogacestat. Program activity was quantified as in **Figure 4D**, and inhibitor effects were calculated relative to matched N3EFNB + DMSO controls. Posterior distributions were estimated using Bayesian mixed-effects models with cell line as a random intercept.

## SUPPLEMENTARY TABLE LEGENDS

Table S1. Mycoplasma test primers.

Table S2. qPCR primers.

Table S3. Primer SNPs.

Table S4. DEG bulkRNA sequencing (control and N3EFNB conditions).

Table S5. scRNA sequencing ciNMF.

Table S6. scRNA sequencing regulons.

Table S7. siRNA ciNMF additive Bayesian.

Table S8. CAF: N3EFNB treatment Percentage by IF.

S9. CAF: NOTCH Inhibitor Percentage by IF.

Table S10. CAF: NOTCH Ligands Percentage by IF.

Table S11. Fibroblast cell lines: N3EFNB treatment Percentage by IF.

Table S12. Fibroblast cell lines: Co-culture Percentage by IF.

Table S13. Fibroblast cell lines: Co-culture slope.

Table S14. Fibroblast cell lines: Washout Percentage by IF.

Table S15. Fibroblast cell lines: N-1 Percentage by IF.

Table S16. Fibroblast cell lines: N-1 Bayesian Mixed Model: Posterior summary.

Table S17. Fibroblast cell lines: GF Doses Percentage by IF.

Table S18. Fibroblast cell lines: GF Doses. Statistics summary.

Table S19: Significant Phosphosites.

Table S20. ON phosphosites

Table S21. Significant pathways KEGG enrichment analysis.

Table S22. Fibroblast cell lines: Signaling Inhibitors Percentage by IF.

Table S23. Fibroblast cell lines: Signaling Inhibitors Bayesian Mixed Model: Posterior summary.

Table S24. Fibroblast cell lines: NOTCH Inhibitor Percentage by IF.

Table S25. Fibroblast cell lines: NOTCH Ligands Percentage by IF.

## Additional information

### Disclosures

There are no relevant disclosures or conflicts of interest.

## Notes

### Competing Interest Statement

The authors have declared no competing interest.

