## Supplementary Figures for "Tumour-stroma signaling drives a reversible program with neural-like features in cancer-associated fibroblasts"

Supplementary Fig. 1

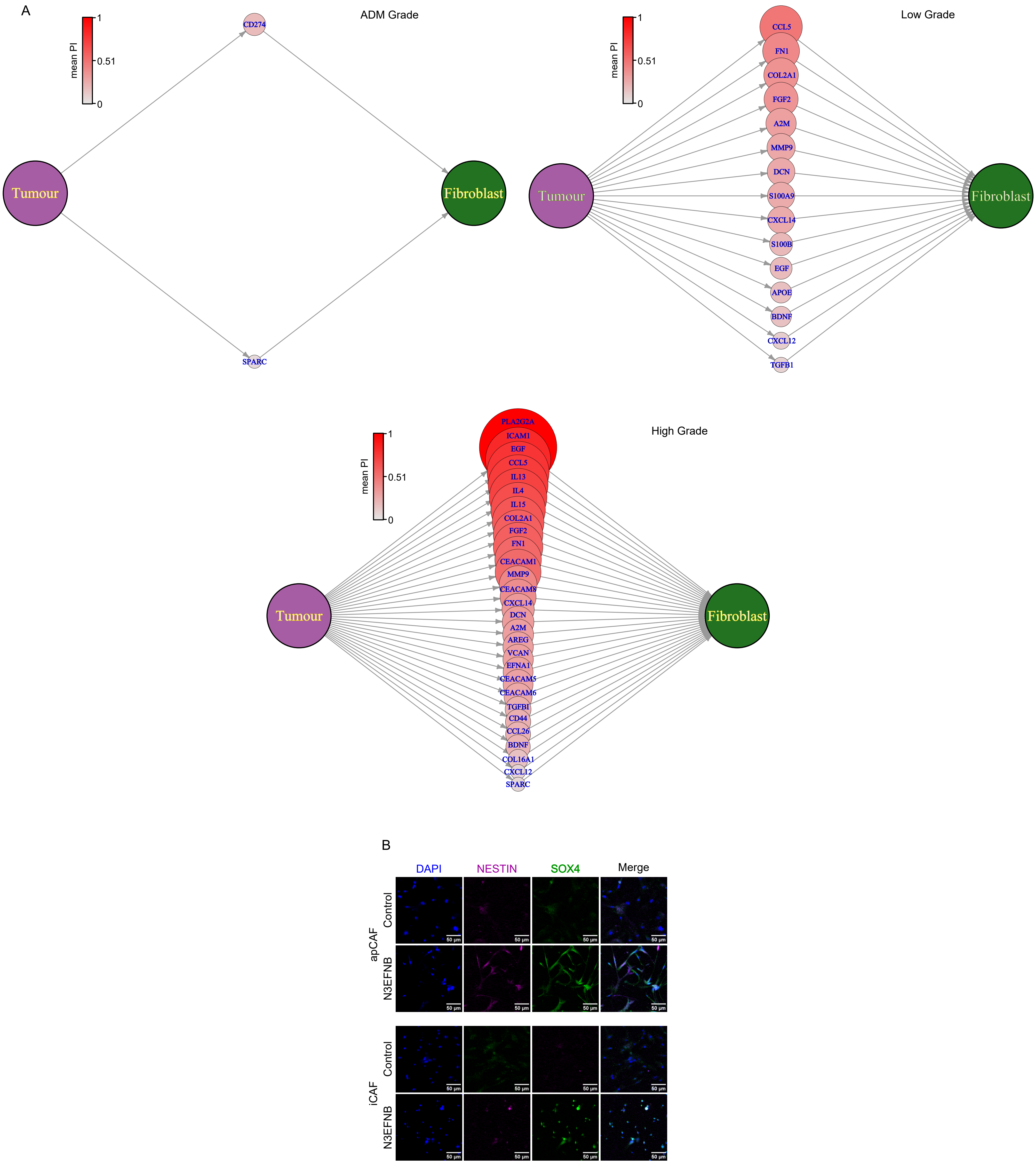

### Supplementary Fig. 2

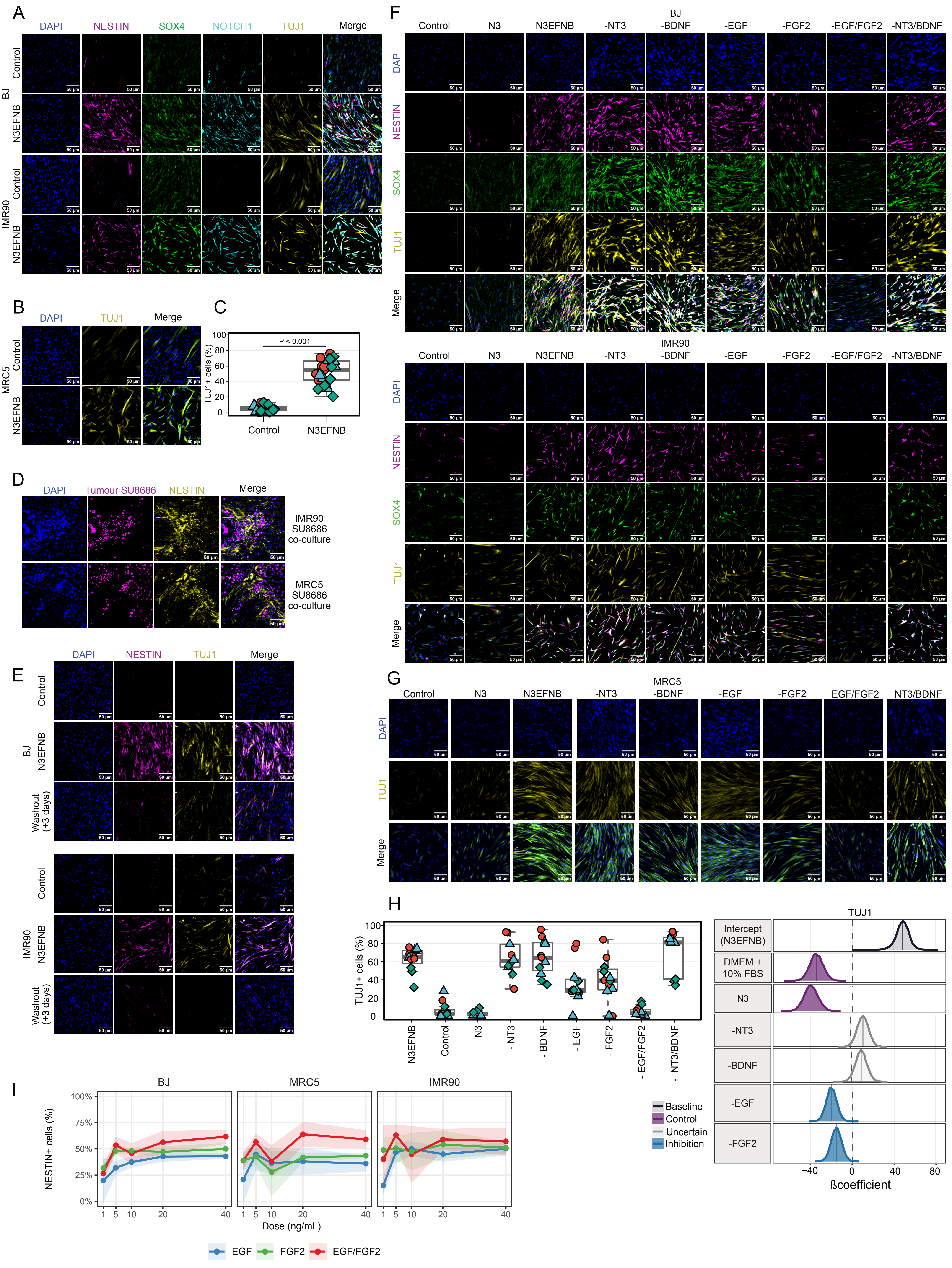

Supplementary Fig. 3

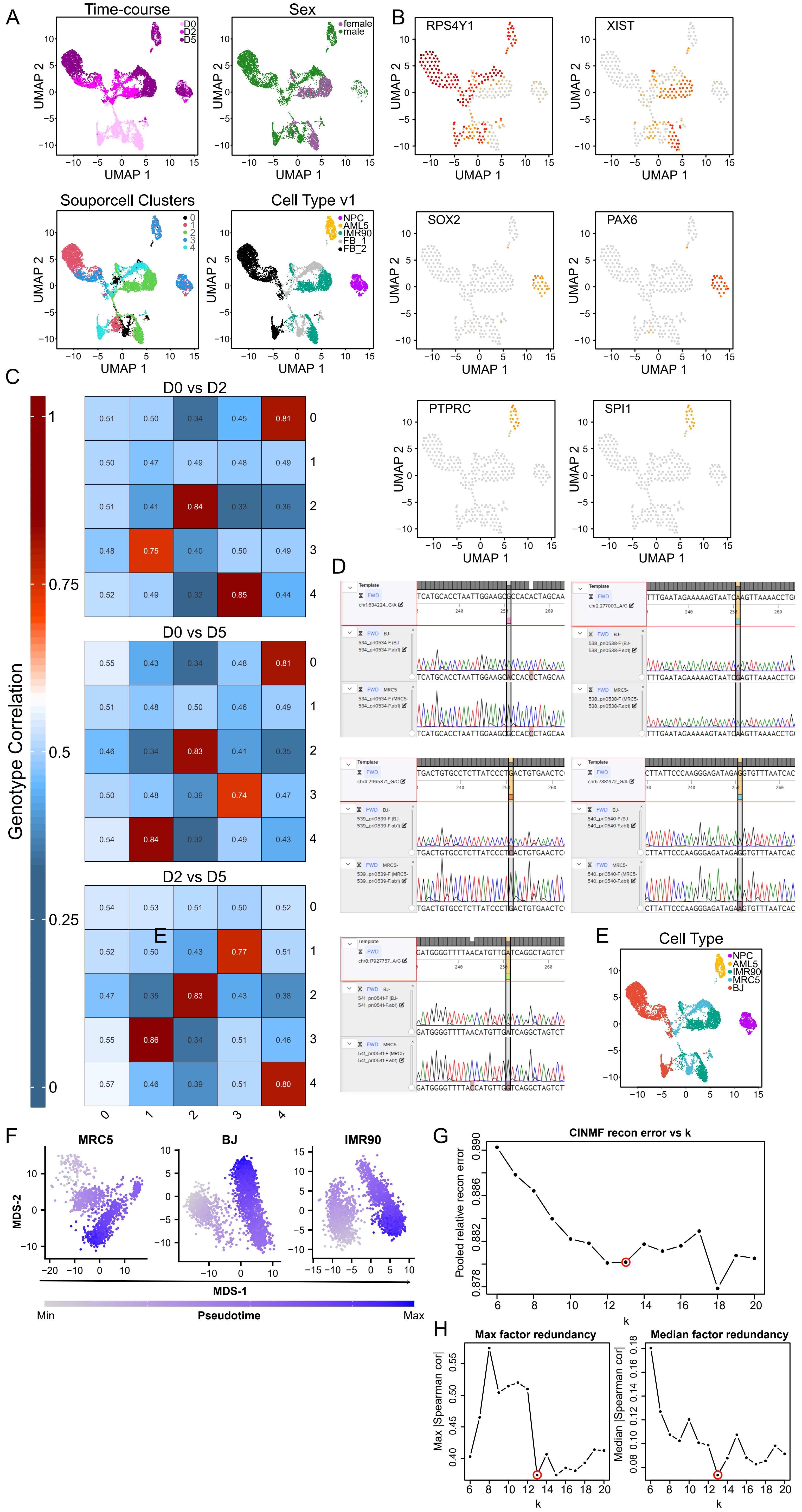

Supplementary Fig. 4

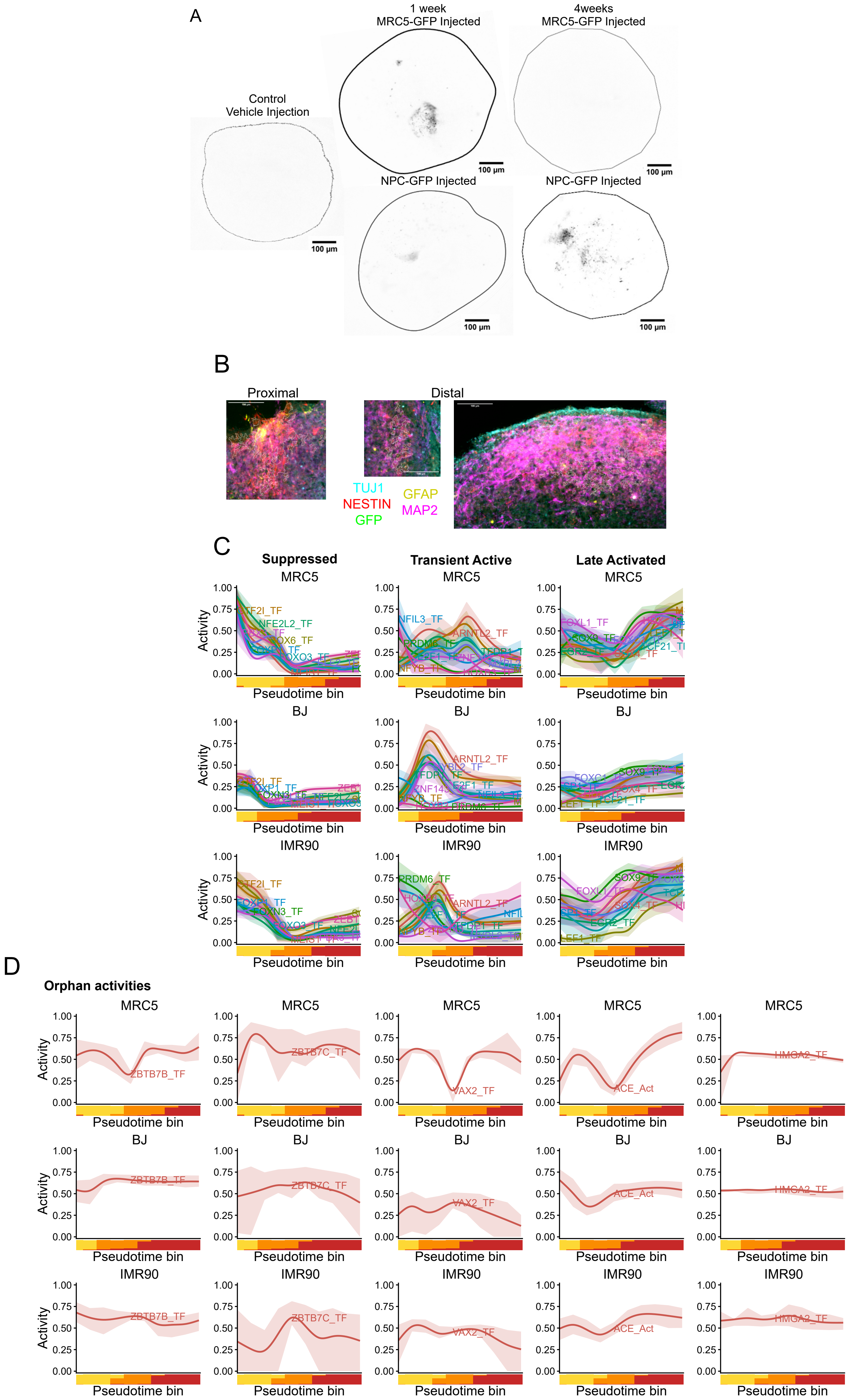

Supplementary Fig. 5

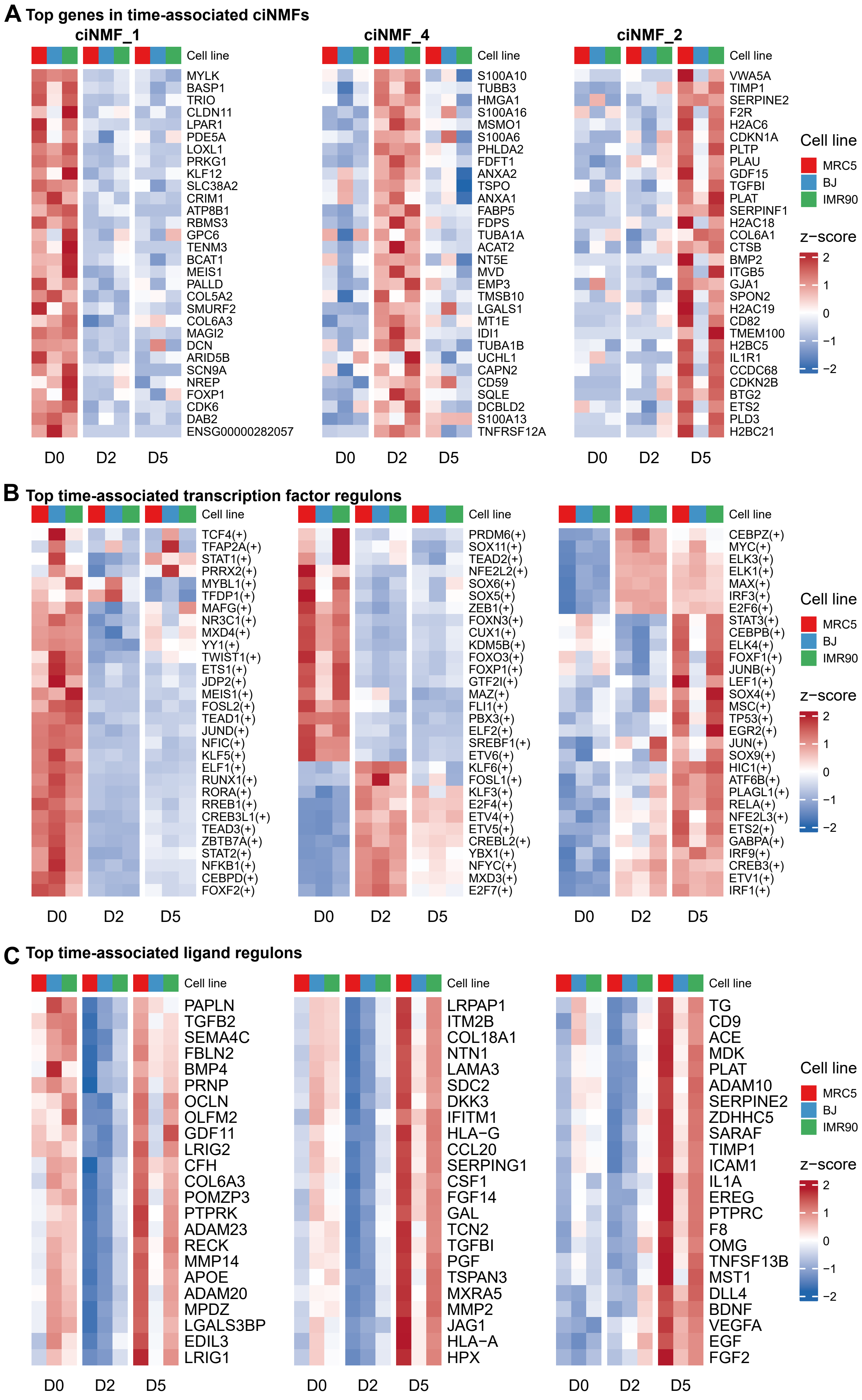

Supplementary Fig. 6

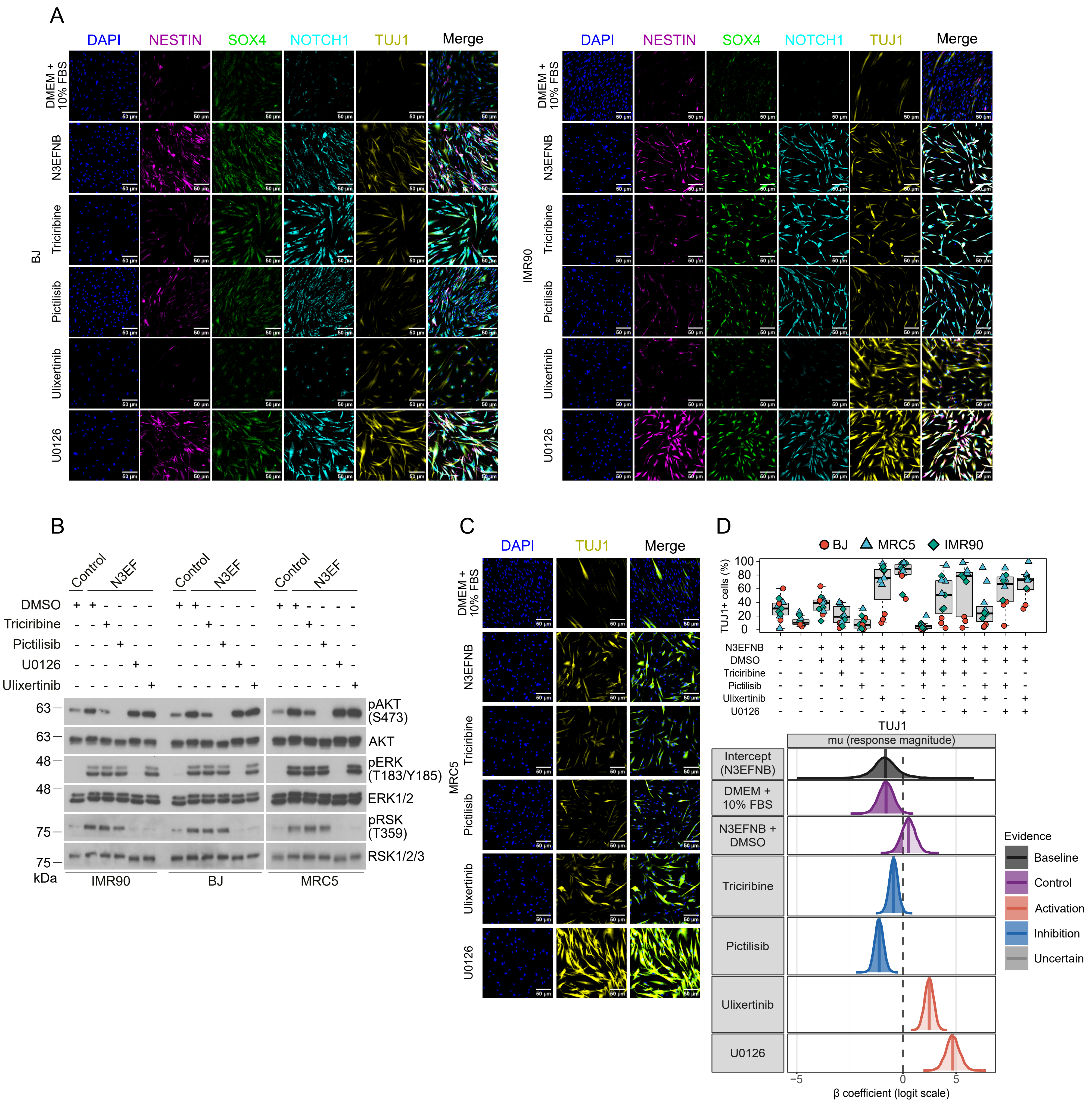

Supplementary Fig. 7

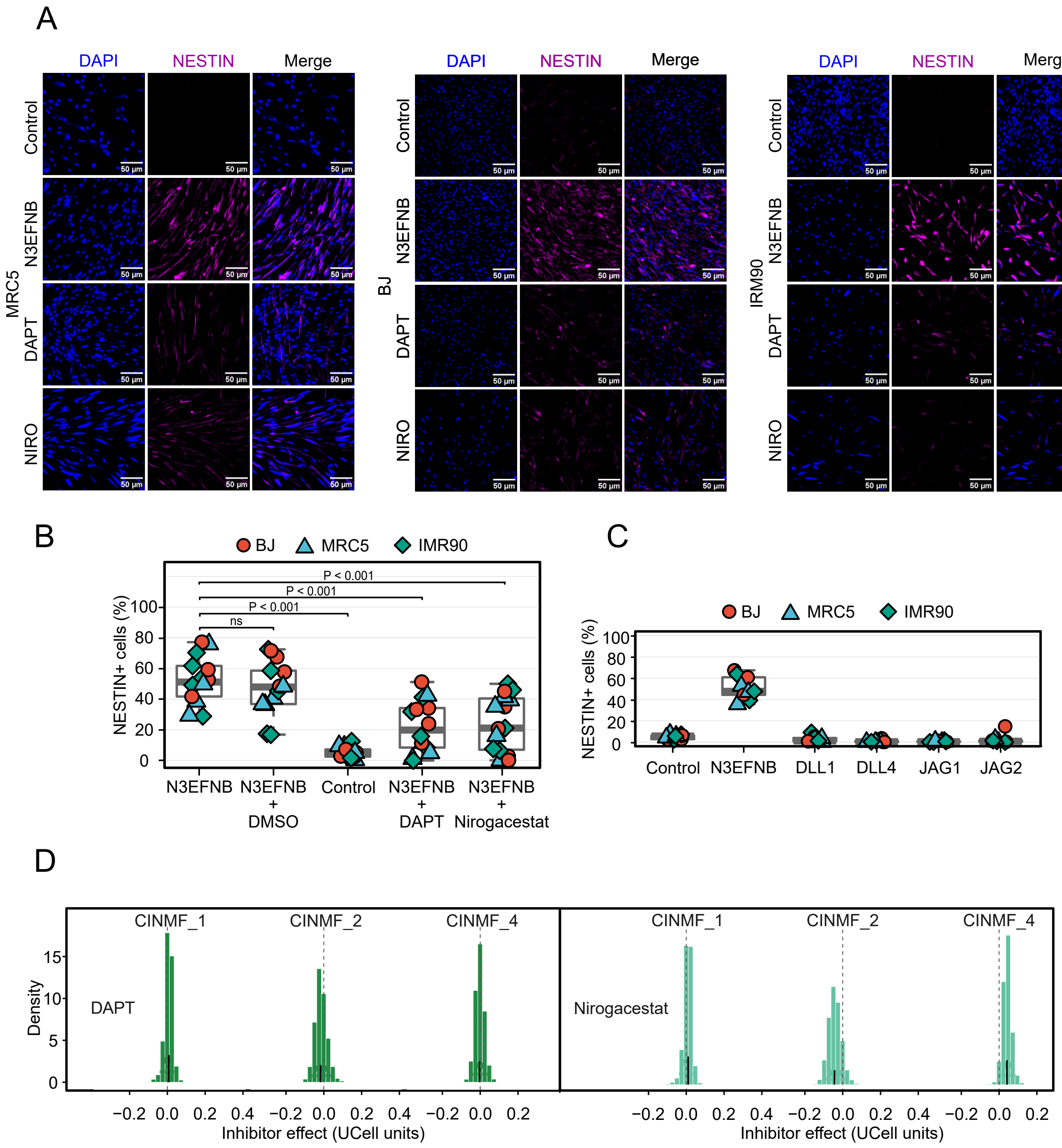
